# Pharmacological SOS1 inhibitor BI-3406 demonstrates *in vivo* anti-tumor activity comparable to SOS1 genetic ablation in KRAS mutant tumors

**DOI:** 10.1101/2024.09.18.613686

**Authors:** Fernando C. Baltanas, Maximilian Kramer-Drauberg, Rósula Garcia-Navas, Enrico Patrucco, Ettore Petrini, Heribert Arnhof, Andrea Olarte-San Juan, Pablo Rodríguez-Ramos, Javier Borrajo, Nuria Calzada, Esther Castellano, Barbara Mair, Kaja Kostyrko, Marco H. Hofmann, Chiara Ambrogio, Eugenio Santos

**Affiliations:** Laboratorio 1. Centro de Investigación del Cáncer, Instituto de Biología Molecular y Celular del Cáncer, CSIC-Universidad de Salamanca and CIBERONC, 37007 Salamanca, Spain; Instituto de Biomedicina de Sevilla, IBiS/Hospital Universitario Virgen del Rocío/CSIC/Universidad de Sevilla and Departamento de Fisiología Medica y Biofísica, Universidad de Sevilla, Sevilla, Spain; Department of Molecular Biotechnology and Health Sciences, Molecular Biotechnology Center, University of Torino, Torino, Italy; Departamento de Ciencias Biomédicas y del Diagnóstico. Universidad de Salamanca, 37007 Salamanca, Spain; Laboratorio 5. Centro de Investigación del Cáncer, Instituto de Biología Molecular y Celular del Cáncer, CSIC-Universidad de Salamanca, 37007 Salamanca, Spain; Boehringer Ingelheim RCV GmbH & Co KG, Vienna, Austria

**Keywords:** SOS1, SOS2, GEF, KRAS, LUAD, TME, mouse KO models, SOS1 inhibitors, therapeutic target, BI-3406

## Abstract

Resistance to KRAS^mut^ inhibitors frequently arises, warranting further searches for anti-RAS cancer therapies. We evaluated the tolerability and efficacy of SOS1 pharmacological inhibition in comparison to genetic ablation in different KRAS-dependent tumor settings. Contrary to the rapid lethality caused by SOS1 genetic ablation in SOS2KO mice, SOS1 pharmacological inhibition by its specific inhibitor BI-3406 did not significantly affect animal weight/viability nor cause noteworthy systemic toxicity. In BI-3406-treated KRAS^mut^ MEFs, we observed significantly reduced RAS-GTP levels and RAS downstream signaling, as well as decreased tumor burden and slower disease progression resulting from tumor-intrinsic and extrinsic therapeutic drug effects. In vivo analyses of KRAS^G12D^ allografts in immunocompromised mice and KRAS^G12D^-driven lung adenocarcinomas in immunocompetent mice showed that systemic BI-3406 treatment impaired tumor growth and downmodulated components of the tumor microenvironment comparably to the KRAS^G12D^ inhibitor MRTX1133. Markedly stronger synergistic antitumor effects were observed upon concomitant BI-3406+MRTX113 treatment, confirming SOS1 as an actionable therapy target in RAS-dependent cancers.

## INTRODUCTION

After decades of research, the first direct inhibitors of KRAS G12C oncogenic mutants were recently approved, changing the treatment landscape for patients carrying tumors with this KRAS allele ^1–4^. Unfortunately, rapid appearance of intrinsic and acquired resistance due to multiple potential mechanisms was reported ^5–8^. Within the MAPK pathway, besides secondary KRAS mutations, resistance was shown to be driven by receptor tyrosine kinase upregulation or activation of the wild-type (WT) RAS isoforms ^9–11^. To enhance durability of response the quest for new anti-RAS cancer therapies is continuing with a focus on KRAS inhibitor combinations with drugs targeting downstream or upstream components of RAS pathway ^12–16^.

The ubiquitously expressed members of the SOS Guanine Exchange Factor (GEF) family (SOS1 and SOS2) are the most universal and functionally relevant activators of RAS proteins in a variety of biological contexts ^12,13,17,18^. Initial analyses of constitutive SOS1 and SOS2 knockout (KO) mouse strains showed that SOS1 is essential for embryonic development ^19^ but SOS2 is dispensable for reaching adulthood in mice ^20^. Single SOS1 or SOS2 KO is perfectly viable in adult mice, but concomitant SOS1 and SOS2 ablation leads to precipitous death of the animals ^21^. While these observations at the organismal level pointed to some partial functional redundancy between SOS1 and SOS2, subsequent detailed analyses at the cellular level demonstrated the functional prevalence of SOS1 over SOS2 for cellular proliferation and viability in most contexts ^21–23^. Specific functional roles of SOS1 or SOS2 have been observed in different physiological cellular conditions ^24–26^. In pre-clinical mouse models of cancer, SOS1, but not SOS2, is critically required for the development of DMBA/TPA-induced skin tumors ^27^, BCR/ABL-driven chronic myeloid leukemia ^28,29^, or KRAS^G12D^-driven lung adenocarcinoma (LUAD) ^30^. Moreover, the therapeutic efficacy of targeting RAS/MAPK signaling has also been shown in various preclinical settings by using genetic inactivation or pharmacological inhibition ^12,31–33^ further supporting the consideration of SOS1 as a relevant therapeutic target in RAS-dependent tumors.

In this regard, pharmacological inhibitors have been developed in recent years with proven ability to directly block SOS1::RAS interactions ^14,34–38^ or to promote degradation of intracellular SOS1 protein ^39–41^. BI-3406 is an orally bioavailable, potent, and selective SOS1 inhibitor (SOS1i), which has no effect on SOS2 ^35^. It binds to the catalytic domain of SOS1, thus preventing the interaction with GDP-loaded RAS and exchange to GTP-loaded RAS. In SOS2-proficient cells, treatment with BI-3406 results in approximately 50% MAPK pathway downregulation. In RAS-dependent preclinical models, BI-3406 leads to strongest *in vitro* and *in vivo* antitumor activity in combination with other inhibitors of RAS signaling, such as KRAS^G12C^ inhibitors or MEK inhibitors ^6,35,42–44^.

Here, we performed an extensive evaluation of the systemic impact of BI-3406 using our SOS1/2^WT^, SOS1^KO^, SOS2^KO^, or SOS1/2^DKO^ genetic mouse models ^21^. We show that systemic pharmacological inhibition of SOS1 with BI-3406 is well tolerated in SOS1/2^WT^ mice. Moreover, unlike SOS1 genetic ablation *in vivo*, BI-3406 shows no deleterious effects in SOS2-null animals. We also compared the impact of BI-3406 with SOS1^KO^ using *in vitro* and *in vivo* models of KRAS mutant cancer. We demonstrate that SOS1i has an anti-proliferative effect in these models that is similar to the effect of SOS1 genetic ablation we have recently described ^30^. We also find that this is accompanied by downmodulation of RAS downstream signaling in the tumor cells and also results in extensive reprograming of the tumor microenvironment (TME). Previous studies have demonstrated that SOS1 inhibition, in combination with indirect ^35,43,45^ or direct ^42^ inhibitors of RAS signaling, leads to enhanced anti-proliferative effects in KRAS-driven tumors. Here, we also show that a similar effect can be achieved when combining SOS1i with a new class of allele-specific KRAS inhibitors - KRAS G12D targeting drugs. Overall, these observations suggest that SOS1 inhibitors may be good universal combination partners for multiple RAS-targeting strategies.

## RESULTS

### BI-3406 treatment results in negligible systemic effects in healthy naïve or SOS2^KO^ mice

Consistent with our previous characterization of constitutive SOS2^KO^ mice and tamoxifen (TAM)-inducible SOS1^KO^ (Cre^ERT2^; SOS1^fl/fl^) mice ^21^, single SOS1 or SOS2 ablation did not affect the viability of adult mice, but double KO of SOS1 and SOS2 (SOS1/2^DKO^) in adult mice resulted in quick death after only about 2 weeks of TAM treatment (**Fig. 1A**). In contrast to concomitant SOS1/2 genetic ablation, continuous oral administration of BI-3406 (50 mg/kg, bid) in adult mice for 26 days did not affect the survival rate nor produce any noticeable external phenotypic changes in SOS1/2^WT^ animals or in constitutive-null SOS2^KO^ mice (**Fig. 1B**).

**Fig. 1.**
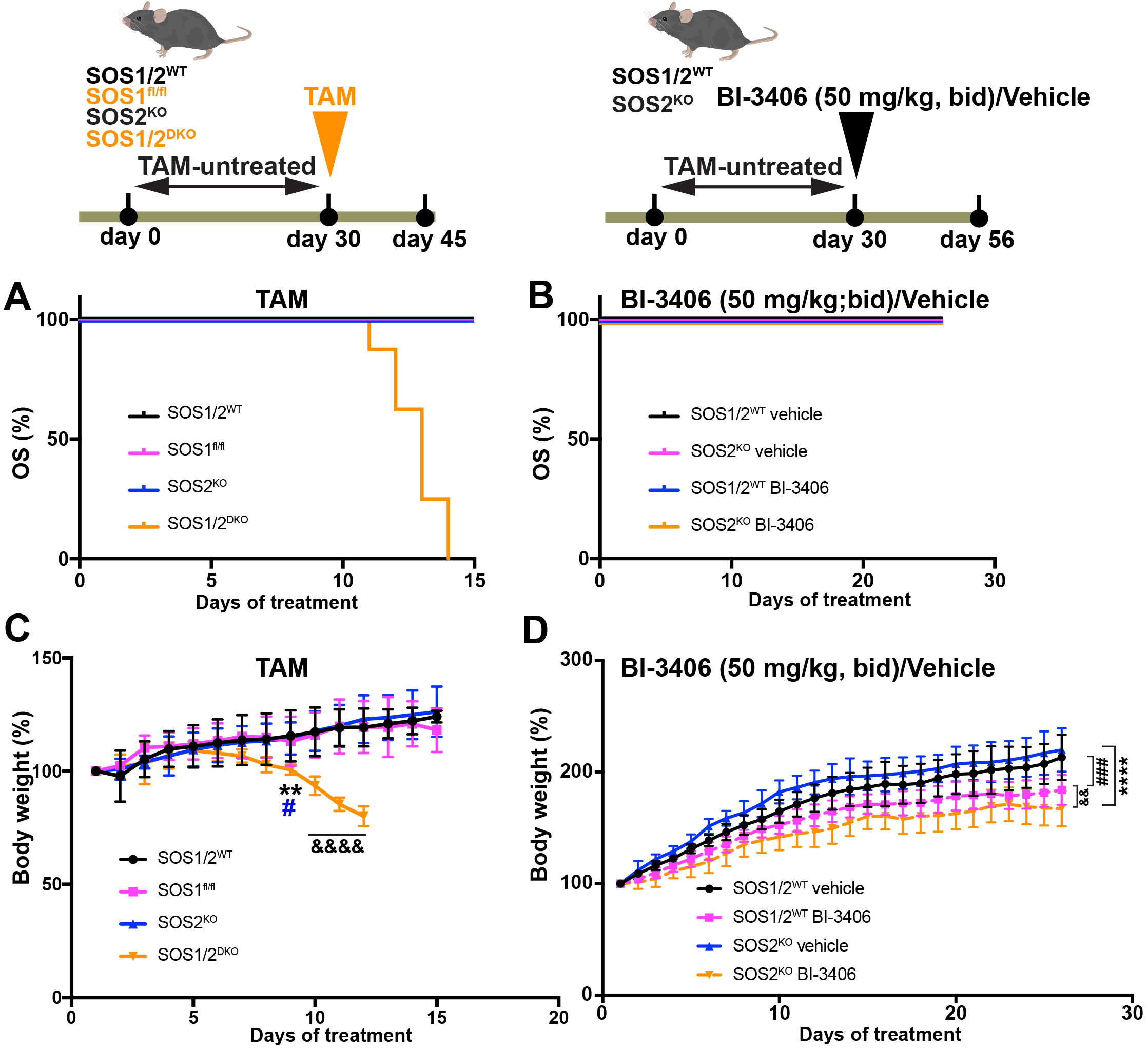
Assessment of *in vivo* toxicities resulting from genetic ablation or pharmacological inhibition of SOS1 in mice. **(A)** Kaplan-Meier survival plot of TAM-induced (TAM-containing chow diet starting at 1 month of age) SOS1/2^WT^, SOS1^fl/fl^, SOS2^KO^ (black, pink and blue lines, respectively) and SOS1/2^DKO^ mice (orange line). **(B)** Kaplan-Meier survival plot of SOS1/2^WT^ or SOS2^KO^ experimental groups orally treated with vehicle (Natrosol; via gavage) or BI-3406 (50 mg/kg bid; via gavage) for 26 days. **(C)** Body weight measurements at various timepoints (days 1 to 15) during TAM treatment (starting at 1 month of age) in SOS1/2^WT^, SOS1^fl/fl^, SOS2^KO^ and SOS1/2^DKO^ experimental groups. **p< 0.01 vs SOS1/2^WT^; #p<0.05 vs SOS2^KO^; &&&&p< 0.0001 vs rest of groups. Data expressed as mean ± SD. Multiple t-test was used. **(D)** Body weight measurements during BI-3406 or vehicle administration in SOS1/2^WT^ or SOS2^KO^ groups. ###p< 0.001 SOS1/2^WT^ (vehicle-treated) vs SOS1/2^WT^ (BI-3406-treated); ****p< 0.0001 SOS2^KO^ (vehicle-treated) vs SOS2^KO^ (BI-3406-treated); &&p< 0.01 SOS1/2^WT^ (BI-3406-treated) vs SOS2^KO^ (BI-3406-treated). Data expressed as mean ± SD. Multiple t-test was used. For TAM treatments: SOS1/2^WT^ (n=7); SOS1^KO^ (n=9); SOS2^KO^ (n=8); SOS1/2^DKO^ (n=8). Following vehicle (Natrosol) treatment: SOS1/2^WT^ (n=10); SOS2^KO^ (n=8). Following BI-3406 treatment: SOS1/2^WT^ (n=21); SOS2^KO^ (n=30).

We also carefully monitored growth and body weight of 1-month-old mice of the relevant SOS genotypes during genetic or pharmacological disruption of SOS1 (**Fig. 1C, D**). Consistent with our previous observations ^21^ the pattern of body weight progression for the genetically-ablated, single-null SOS2^KO^ or the TAM-induced SOS1^KO^ mice was virtually identical to that of SOS1/2^WT^ mice, whereas SOS1/2^DKO^ mice exhibited a dramatic loss of body weight in comparison to the remaining experimental groups (**Fig. 1C**). In an independent experiment, control (vehicle-treated) SOS1/2^WT^ and control SOS2^KO^ mice showed comparable body weight gain (**Fig. 1D**).

We next compared various serum biochemical parameters between the different experimental groups (**Table 1**). Consistent with our previous reports ^21^ genetic ablation of SOS1/2 resulted in significantly reduced levels of serum albumin and total protein content, elevated levels of various liver-related enzymatic activities and altered levels of triglycerides, total cholesterol, LDH and uric acid (**Table 1A**). In contrast, BI-3406-mediated SOS1 inhibition showed no significant changes in these parameters in SOS1/2^WT^ mice (**Table 1B**) and in SOS2^KO^ (SOS1^WT^/SOS2^KO^) mice produced only a slight increase of GOT, CPK and LDH serum levels (**Table 1B**). Furthermore, the values of the serum-level parameters in BI-3406-treated SOS2^KO^ mice were much closer to those of TAM-treated SOS1/2^WT^ mice than to those of TAM-treated SOS1/2^DKO^ mice (**Table 1**), thus underscoring the low impact of BI-3406-mediated administration in SOS2^KO^ mice at the organismal level. Finally, it should be noted that oral administration of the vehicle (natrosol) in the SOS2^KO^ group did not have any effect on the analyzed biochemical parameters (**Suppl Table 1**), indicating that the mild alterations observed in serum parameters of the BI-3406-treated SOS2^KO^ mice were specifically due to the biological action of BI-3406.

**Table 1.** Blood serum parameters were assessed at day 14 after TAM-treatment **(A)** in SOS1/2^WT^ (n=8), SOS1^KO^ (n=9), SOS2^KO^ (n=10) and SOS1/2^DKO^ (n=8) experimental groups or at day 26 after BI-3406 oral administration (50 mg/kg, bid). **(B)** in SOS1/2^WT^ (n=21) or SOS2^KO^ mice (n=30). *p< 0.05, **p<0.01 vs TAM-treated SOS1/2^WT^. &p< 0.05 vs TAM-treated SOS1^KO^; #p< 0.05, ##p< 0.01 vs TAM-treated SOS2^KO^ ###p< 0.001 vs TAM-treated SOS2^KO^; %%p< 0.01, %%%p< 0.001 vs rest of TAM-treated genotypes; $$ p< 0.01, $$$$p< 0.0001 vs BI-3406-treated SOS1/2^WT^ mice. Data expressed as Mean±SD. One-way ANOVA and Tukey’s correction was used in A and unpaired t-test was used in B. **Alb**: Albumin; **ALP**: Alkaline Phosphatase; **BUN**: Blood urea nitrogen; **Ca**: Calcium; **CPK**: Creatine Phosphokinase; **GGT**: gamma-glutamyl transferase; **Glc**: Glucose; **GOT**: glutamic oxaloacetic transaminase; **GPT**: Glutamate-Pyruvate Transaminase; **LDH**: lactate dehydrogenase; **T-Bil**: Total bilirubin; **T-Chol**: Total Cholesterol; **TG**: Triglyceride; **T-Prot**: Total Protein; **UA**: **Uric acid**.

We also analyzed the impact of BI-3406 treatment on hemogram parameters in our SOS1/2^WT^ and SOS2^KO^ mouse strains at the beginning (0 d), mid-term (13 d) and the end (26 d) of treatment (**Table 2**). Comparison of these parameters between SOS1/2^WT^ and SOS2^KO^ animals treated with BI-3406 revealed no significant differences at day 13 but at day 26 several slight changes were observed (**Table 2**). A slight, although statistically significant, reduction of lymphocytes was observed, likely resulting from an increased percentage of neutrophils, as well as an increase in the total number of large unstained cells and a slight alteration in erythrocyte-related parameters such as hemoglobin, hematocrit, and red cell distribution width (**Table 2**). Importantly, comparison of these parameters in BI-3406-treated SOS2^KO^ mice with those previously measured in SOS1/2^DKO^ mice ^21^ showed that the alterations of hematological parameters measured in BI-3406-treated SOS2^KO^ mice were quantitatively milder than those measured following combined genetic depletion of SOS1 and SOS2.

**Table 2.** Hematological parameters from peripheral blood were assessed previous (0d) and at 13 and 26 days after initiation of BI3406 oral administration in SOS1/2^WT^ (n=12) and SOS2^KO^ (n=12) experimental groups. *p< 0.05, **p< 0.01 vs BI3406-treated SOS1/2^WT^ mice (50 mg/kg, bid; 26d). Data expressed as Mean±SD. **CH**: mean optical hemoglobin content of intact RBCs; **CHCM**: cellular hemoglobin concentration mean; **HCT**: Hematocrit; **HGB**: Hemoglobin; HDW: **LUC**: large unstained cells; **MCH**: mean corpuscular hemoglobin; **MCHC**: mean corpuscular hemoglobin concentration; **MCV**: Mean corpuscular volume; **MPV**: Medium Platelet Volume. **PLT**: platelet; **RBC**: red blood cells; **RDW**: red cell distribution width; **WBC**: white blood cells.

The almost complete absence of systemic effects of BI-3406-mediated SOS1 inhibition was further confirmed by comparing histological preparations of different tissues obtained from necropsies of SOS1/2^WT^ or SOS2^KO^ mice after TAM-treatment or oral administration of BI-3406 (**Fig. 2**). Consistent with our previous reports ^21^, SOS1/2^DKO^ mice exhibited obvious histological defects in the thymus, spleen and liver as compared to SOS1/2^WT^ controls, whereas other organs such as the kidney, heart or lung did not display any visible morphological alterations (**Fig. 2**, left panels). These results were in concordance with the observed reduction in lymphocytes and the increase of transaminases (**Table 1A**). In contrast, SOS1/2^WT^ mice or SOS2^KO^ mice treated with BI-3406 did not show obvious histopathological findings (**Fig. 2**, right panels).

**Fig. 2.**
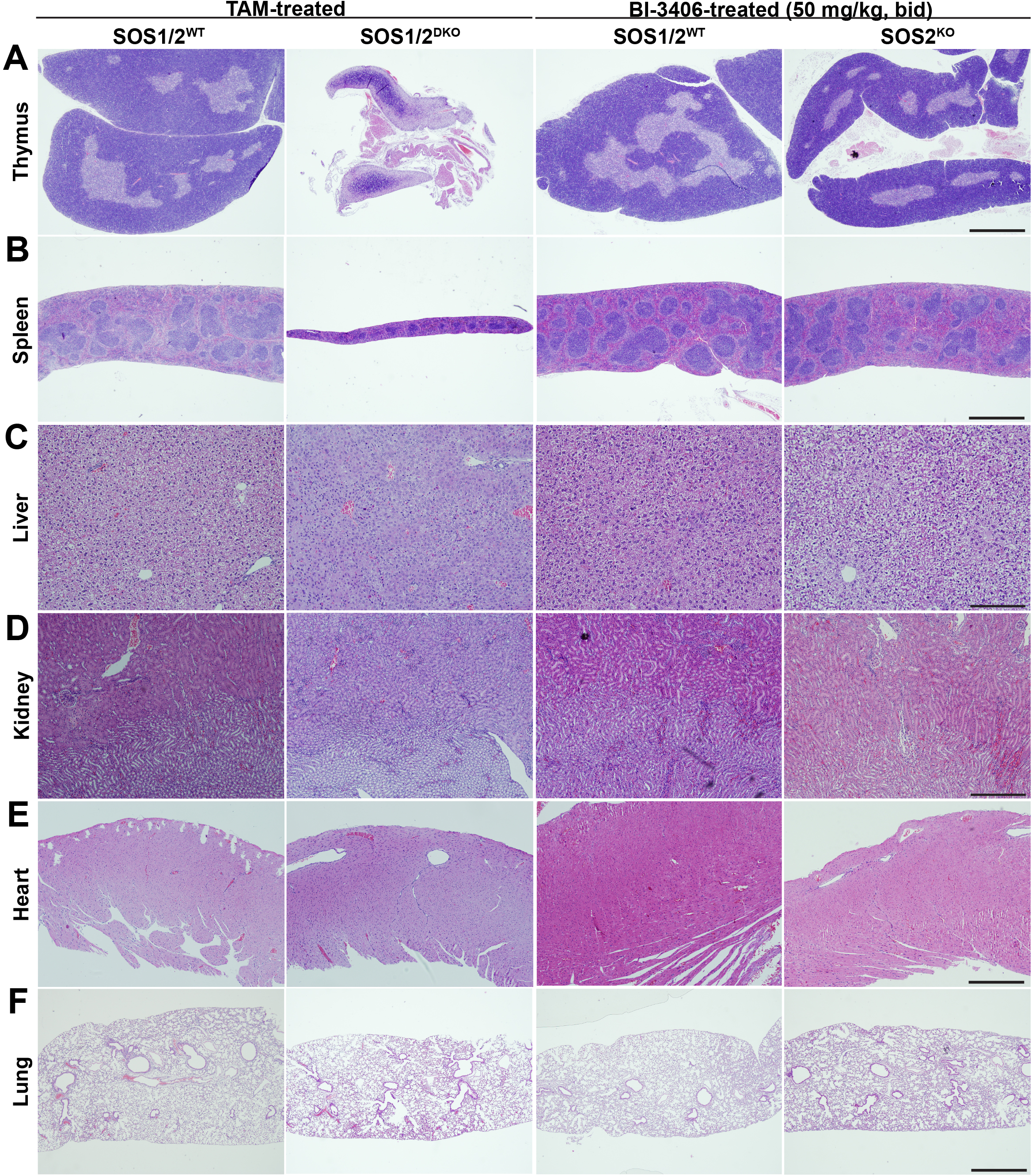
Histological examination of the impact of SOS1 genetic or pharmacological depletion in naïve and SOS2^KO^ mice. Representative images of paraffin-embedded sections stained with H&E from thymus (A), spleen (B), liver (C), kidney (D), heart (E) and lung (F) of TAM-treated (TAM-containing chow diet for 12 days) SOS1/2WT and SOS1/2DKO mice or BI-3406-treated (50 mg/kg, bid, via gavage for 26 days) SOS1/2WT and SOS2KO groups. All treatments started at 1 month of age. Scale bars: (A, B, F): 500 μm; (C-D): 100 μm; (E): 200 μm.

Nevertheless, in depth examination revealed minor structural alterations in the liver and kidney of SOS2^KO^ mice treated with BI-3406 (**Suppl Fig. S1A**). In particular, the liver of BI-3406-treated SOS2^KO^ mice displayed occasional clusters of morphologically altered hepatocytes characterized by smaller and darker nuclei and retracted cytoplasm (**Suppl Fig. S1A**). These morphological alterations might be related to the slight increase in transaminase levels detected in BI-3406-treated SOS2^KO^ mice (**Table 1B**). These findings were not observed in the SOS1/2^WT^ BI-3406 treated animals (**Suppl Fig. S1A**). In addition, we detected structural disorganization of the epithelial cells covering the surface of the proximal and distal tubules of the kidneys of BI-3406-treated SOS2^KO^ mice (**Suppl Fig. S1A**), although the nephritic glomeruli did not show any observable morphological alterations (**Suppl Fig. S1A**). These kidney alterations might correlate with the minor increase in CPK levels detected in the serum of BI-3406-treated SOS2^KO^ mice (**Table 1B**). Detailed examination of various other organs from these mice, including the pancreas (**Suppl Fig. S1A**), did not uncover any other morphological alterations that could be linked to BI-3406 treatment of SOS2^KO^ mice. In addition, the vehicle control (natrosol; **Suppl Fig. S1B**) did not show any gross structural alterations in any of the analyzed organs. A detailed histopathologic scoring of the morphological studies performed in different tissues and organs of our BI-3406-treated or untreated wild-type animals or animals with genetic ablation of SOS1 and/or SOS2 is shown in Supplementary Table 2.

Overall, all the above observations demonstrate that the effects resulting from *in vivo* pharmacological SOS1 inhibition in SOS2^KO^ mice were significantly milder than those observed in the SOS1/2^DKO^ mice generated by genetic ablation of SOS1 in SOS2^KO^ mice.

### Genetic ablation and pharmacological inhibition of SOS1 result in similar functional and mechanistic effects in KRAS^mut^ cells

Following the tolerability experiments, we proceeded to evaluate the functional impact of BI-3406 treatment in cells driven by different KRAS mutants. For this purpose, we evaluated the effects of BI-3406-mediated inhibition of SOS1 *versus* genetic SOS1 deletion using immortalized MEFs of the relevant SOS genotypes (SOS1/2^WT^, SOS1^fl/fl^, SOS2^KO^, SOS1/2^DKO^) ^22^ expressing several common mutants of human KRAS (G12C, G12D and G12V) (**Suppl Fig. S2A)**.

We first assessed the impact of BI-3406 treatment on the viability of SOS/KRAS^mut^ MEFs by performing drug response assays using a 3-day BI-3406 treatment, which, in the SOS1/2^WT^ genetic background, resulted in a dose-dependent reduction of viability in G12C, G12D, and G12V KRAS^mut^-driven cells (**Fig. 3A**). Genetic SOS2 depletion further sensitized KRAS^mut^ cells to BI-3406 treatment, leading to a stronger anti-proliferative effect, likely due to complete loss of SOS GEF activity. As expected, SOS1 genetic deletion abrogated the sensitivity to BI-3406, confirming drug selectivity for SOS1 and suggesting the lack of off-target effects of BI-3406 at the cellular level.

**Fig. 3.**
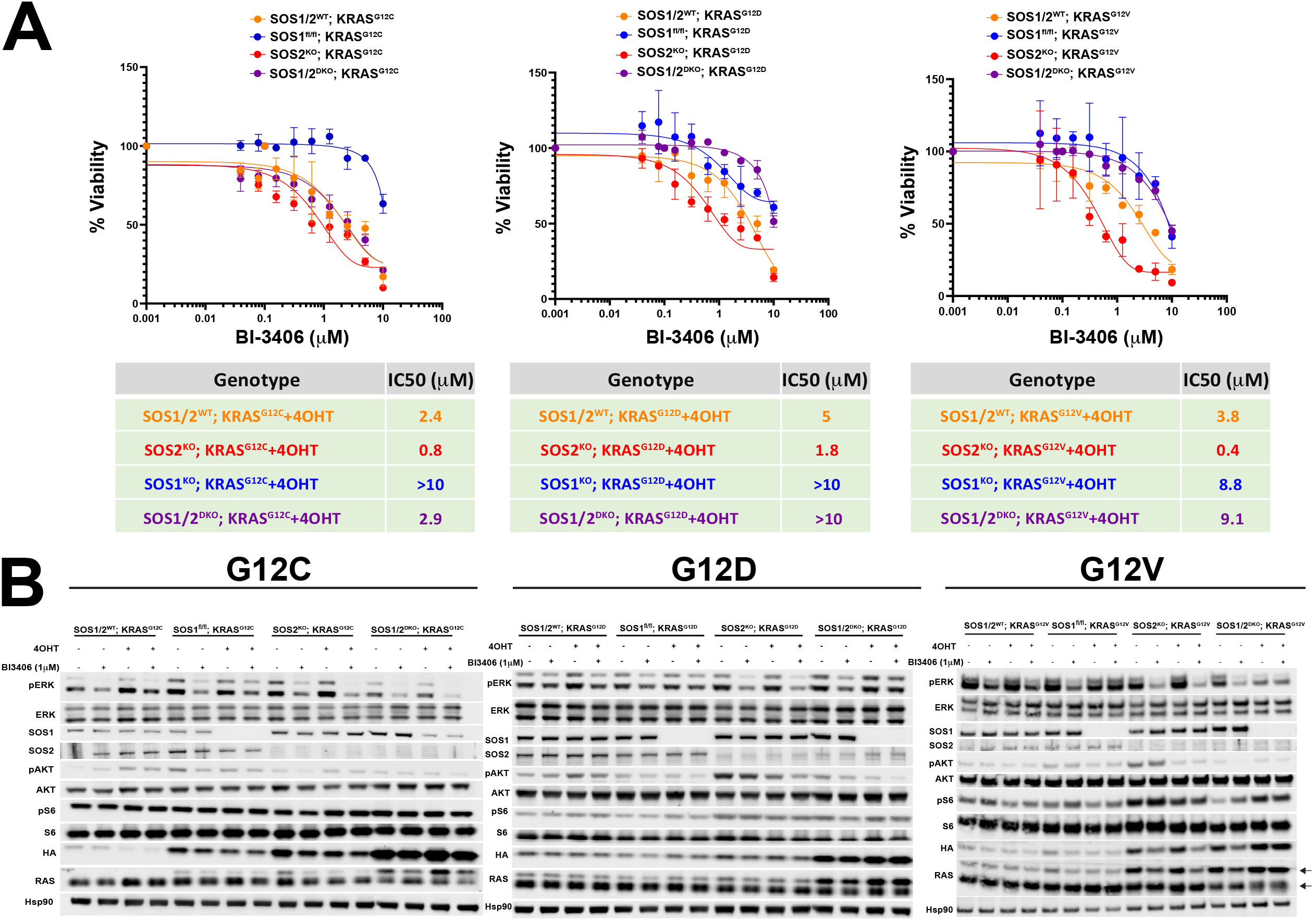
Determination of BI-3406 sensitivity in oncogenically KRAS mutated immortalized MEFs. **(A)** Dose-response curves to BI-3406 in immortalized MEFs of the four SOS genotypes (SOS1/2^WT^, SOS1^KO^, SOS2^KO^ and SOS1/2^DKO^) expressing exogenous HA-tagged KRAS^G12C^, KRAS^G12D^ or KRAS^G12V^ in presence of 4OHT (0.3 μM). Data are represented as mean ± SD. n=3 independent samples per experimental group. Results are representative of one of three similar experiments. **(B)** Representative Western-blot images of RAS downstream signaling factors in SOS1/2^WT^, SOS1^KO^, SOS2^KO^ and SOS1/2^DKO^ MEFs expressing oncogenic KRAS mutations (G12C, G12D, G12V) and treated or untreated with 4OHT and/or BI-3406 (individually or combined). n=3 independent samples per experimental group. Arrows indicate exogenous HA-tagged (upper) and endogenous WT RAS (lower).

To further investigate the impact of SOS1 disruption on KRAS^mut^ cells, we performed live cell imaging-based proliferation assays of KRAS^mut^ cell lines treated with SOS1i and/or subjected to single or double SOS1 or SOS2 knock-out (**Suppl Fig. S2B-C**). In agreement with the drug response assays, the genetic disruption of SOS1 caused a mild, but significant decrease in the proliferative ability of the immortalized KRAS^mut^ cells (**Suppl Fig. S2B**). Interestingly, pharmacological inhibition of SOS1 exerted a more potent anti-proliferative effect than SOS1^KO^, with a milder phenotype in KRAS^G12D^-driven cells compared to either KRAS^G12C^- or KRAS^G12V^-driven cells (**Suppl Fig. S2C**). Of note, BI-3406 treatment had an enhanced anti-proliferative effect in KRAS^mut^ cells in the absence of SOS2 (SOS2^KO^/KRAS^mut^ cells) (**Suppl Fig. S2C**). As expected, genetic depletion of SOS1 with 4-hydroxtamoxifen (4OHT) treatment, alone or in combination with SOS2^KO^, rendered cells insensitive to BI-3406 treatment.

Immunoblotting analysis of KRAS^mut^ cells revealed a reduction in ERK phosphorylation (pERK) upon BI-3406 treatment across the different genotypes, with a more pronounced effect in KRAS^mut^/SOS2^KO^ cells, again suggesting functional redundancy or a partially compensatory role of SOS2 upon SOS1 inhibition (**Fig. 3B**). Of note, genetic deletion of SOS1 did not show any significant pERK reduction, likely due to prolonged 4OHT exposure (>10 days) allowing for rebound mechanisms to occur before the start of our assays. Our overall comparison of the different KRAS^mut^ cell lines tested pointed to a stronger, more pronounced impairment of ERK activation in the KRAS^G12C^ cell lines than in the other two mutant genotypes (**Fig. 3B**). In contrast, no consistent changes in AKT or S6 phosphorylation levels were observed in any of the genotypes or experimental conditions tested (**Fig. 3B**).

We next evaluated the differential abilities to respond to EGF stimulation displayed by our SOS1/2^WT^ and SOS-less KRAS^mut^ immortalized MEF cell lines after undergoing genetic or pharmacological ablation of SOS1 (**Fig. 4**). Active RAS pull down assays revealed that individual genetic disruption of SOS1 or SOS2 already resulted in sizeable reduction of RAS-GTP formation as compared to SOS1/2^WT^ cells, but only the concomitant ablation of SOS1 and SOS2 resulted in almost complete loss of RAS-GTP upon EGF stimulation (**Fig. 4A**). Consistent with the cell proliferation assays (**Suppl Fig. S2C**), these results also point to the dependency on SOS1 or SOS2 genetic depletion of the KRAS^mut^ cell lines with regards to RAS-GTP formation and activation of RAS downstream signals (pERK, pAKT) in response to external EGF treatment (**Fig. 4A**).

**Fig. 4.**
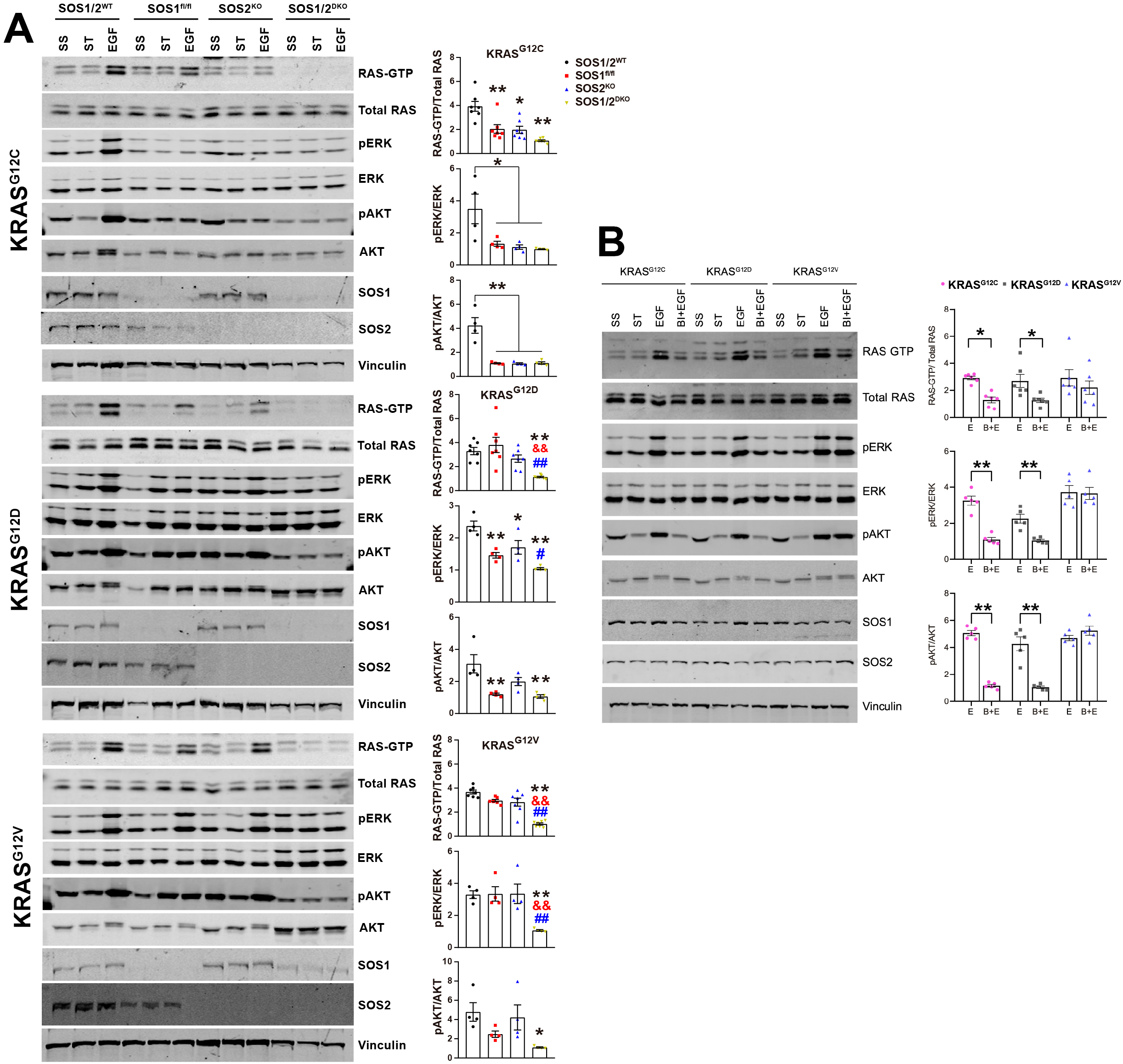
Effect of SOS1 silencing in the activation of RAS/MAPK signaling pathway upon EGF stimulation. **(A)** Immortalized MEFs of SOS1/2^WT^, SOS1^fl/fl^, SOS2^KO^, and SOS1/2^DKO^ genotypes harboring KRAS^G12C^, KRAS^G12D^ or KRAS^G12V^ were 4OHT-treated (0.3 μM) for 12 days, serum-starved (ST) for 24 hours and subsequently stimulated with EGF (100 ng/ml) for 2 minutes. The images are representative Western-blots of active RAS pull down assays (RAS-GTP) and the corresponding signaling of pERK and pAKT, as well as the expression of SOS1 and SOS2 in each case. Vinculin was used as a loading control. The quantitation of RAS/ERK/AKT activation levels (graphs in the right columns) was performed by calculating the ratio of increased RAS-GTP, pERK and pAKT levels compared to ST cells. Data represents mean ± SD (n=5 independent samples per experimental group). *p < 0.05 and **p < 0.01 vs SOS1/2^WT^ group; &&p < 0.01 vs SOS1^KO^; ##p < 0.01 vs SOS2^KO^. One-way ANOVA and Tukey’s test. **(B)** Immortalized MEFs of the SOS1/2^WT^ genotype expressing KRAS^G12C^, KRAS^G12D^ and KRAS^G12V^ that were ST for 24 hours and treated (B+E) or untreated (E) with BI-3406 (1 µM) for 2 hours, and subsequently stimulated with EGF (100ng/ml) for 2 minutes. The images shown are representative Western-blots of RAS pull down activation assays (RAS-GTP) and the corresponding signaling of pERK and pAKT, as well as the expression of SOS1 and SOS2 in each case. Vinculin was used as a loading control. The quantitation of RAS/ERK/AKT activation was performed by calculating the ratio of increased RAS-GTP, pERK and pAKT levels compared to ST cells. Data shown as mean ± SD. (n=6 independent samples per experimental group). *p < 0.05 and **p < 0.01 vs EGF+BI-3406 (B+E)-treated group. One-way ANOVA and Tukey’s test. SS: Steady state, ST: Serum-starved.

Importantly, our evaluation of the impact of BI-3406 on the levels of RAS activation and downstream RAS/MAPK signal transmission in SOS1/2^WT^ immortalized MEFs harboring different human KRAS^mut^ forms also demonstrated that BI-3406 treatment resulted in a stronger reduction of RAS-GTP levels as well as pERK and pAKT levels in KRAS^G12C^- and KRAS^G12D^-expressing MEFs as compared to cells harboring KRAS^G12V^ mutation (**Fig. 4B**). Since only one MEF line was used here for each genotype, further work using different cell lines will be required in future to confirm this observation. In any event, it should be noted that the three oncogenic variants reached comparable levels of SOS1 protein ablation upon 4OHT administration **(Suppl Figure S3A).**

In summary, our results demonstrated that BI-3406-mediated pharmacological inhibition of SOS1 exhibited comparable antiproliferative properties to those observed upon genetically-mediated SOS1 ablation in oncogenic KRAS mutated MEFs.

### Genetic or pharmacological inhibition of SOS1 impairs intrinsic tumor growth and downmodulates extrinsic protumorigenic signals in KRAS^G12C^ and KRAS^G12D^ allografts

We next aimed to compare the *in vivo* impact of genetic (4OHT-induced) versus pharmacological (BI-3406 treatment) inhibition of SOS1 on the growth of mouse allografts of oncogenic KRAS^mut^ MEFs. For this purpose, immunocompromised mice were implanted with either SOS1^fl/fl^/KRAS^G12C^ or SOS1^fl/fl^/KRAS^G12D^ MEFs, and randomized for treatment with vehicle, 4OHT or BI-3406.

In KRAS^G12C^ allografts, genetic depletion of SOS1 mediated by 4OHT administration resulted in significant impairment of tumor growth, as assessed by the markedly reduced area and volume of the tumor allografts in comparison to the vehicle-treated samples (**Figs. 5A-B** and **Suppl Fig. S4A**). In this model, pharmacological inhibition of SOS1 mediated by BI-3406 treatment also resulted in a stronger anti-tumor effect than genetic ablation with 4OHT in SOS1^KO^ mice. Of note, the efficiency of SOS1 ablation following *in vivo* 4OHT administration was lower (∼50%) in comparison to 4OHT-treated oncogenic MEFs (∼80%) (**Suppl Fig S3**). Characterization of the tumor allograft explants from these assays by means of immunohistochemistry (**Suppl Fig. S4**) or by immunoblotting (**Suppl Fig. S5A**) also showed a direct correlation of the observed anti-tumor effect of genetic or pharmacological ablation of SOS1 with the reduced levels of cell proliferation and ERK phosphorylation detected in the corresponding tumor explants (**Fig. 5B** and **Suppl Fig. S4B-C**).

**Fig. 5.**
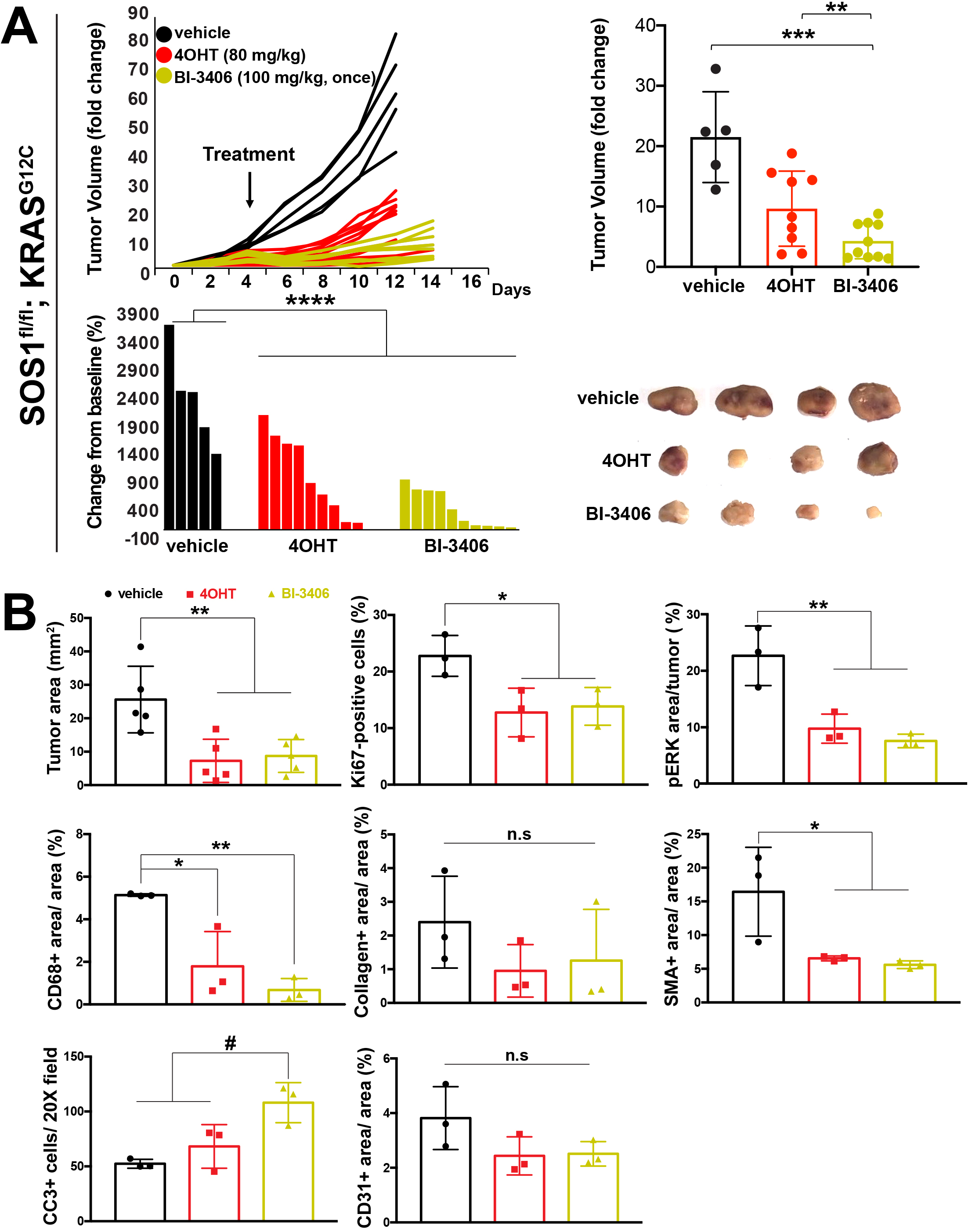
Allograft assays of (genetically or pharmacologically) SOS1-disrupted KRAS^G12C^ MEFs. **(A)** Immortalized SOS1^fl/fl^ MEFs stably expressing exogenous KRAS^G12C^ were injected subcutaneously into nude mice. Once tumors reached a size of 200 mm^3^, animals were treated with vehicle (via gavage), 4OHT (80 mg/kg; intraperitoneally) or BI-3406 (100 mg/kg, once daily; via gavage) for 12 days. The mean fold-change in tumor volume relative to initial tumor volume is shown. The arrow indicates treatment starts. Waterfall plots of individual tumor responses from vehicle, 4OHT or BI-3406-treated are depicted for day 12 and day 14. n=5 (vehicle-treated); n=9 (4OHT-treated); n=10 (BI-3406-treated). Bar graph represents tumor volume fold change relative to treatment start. Error bars represent mean ± SD. **p < 0.01, ***p < 0.001 and ****p < 0.0001. One-way ANOVA was used. The panel includes representative images of SOS1^fl/fl^/KRAS^G12C^ allografts at day 16 after tumor implantation treated (starting at day 4; arrow) with vehicle, 4OHT or BI-3406. **(B)** Quantitation of total tumor area, the percentage of Ki67-positive cells, the percentage of pERK stained area with respect of total tumor area, the percentage of CD68-stained area, the percentage of collagen-stained area, the percentage of SMA-stained area, the number of CC3-positive cells per 20X field and the percentage of CD31-positive area. n=5 tumors per group (for H&E studies) and n=3 tumors per group (for the rest assays). Data shown as mean ± SD. *p < 0.05, **p < 0.01 vs vehicle-treated group; #p < 0.05 vs BI-3406-treated group. One-way ANOVA and Tukey’s test. CC3: cleaved caspase 3; SMA: smooth muscle actin. n.s: not significant.

Furthermore, the immunohistochemical analysis of these allograft explants also clearly showed that, in comparison with vehicle-treated counterparts, both the inhibition of SOS1 by BI-3406 or the 4OHT-induced genetic ablation of SOS1 caused significant downmodulation of pro-tumorigenic components of the tumor microenvironment (TME) (**Fig. 5B** and **Suppl Fig. S4**). In particular, CD68-positive tumor-associated macrophages and SMA-immunoreactive cancer-associated fibroblasts (CAFs) were significantly reduced in the stromal TME of 4OHT-treated and BI-3406-treated allograft explants in comparison with vehicle-treated animals **(Fig. 5B** and **Suppl Fig. S4D-F)**. Cleaved caspase 3 (CC3) immunohistochemical assays of the allograft explants also showed that the BI-3406-treated samples displayed higher levels of cell death than the corresponding 4OHT-treated and vehicle-treated samples **(Fig. 5B** and **Suppl Fig. S4G)**. Finally, as previously described ^30^, CD31 immunostaining demonstrated that SOS1 ablation did not quantitatively impact the intratumoral vasculature **(Fig. 5B** and **Suppl Fig. S4H).**

We next evaluated the *in vivo* consequences of genetic and pharmacological SOS1 inhibition on mouse allografts harboring the oncogenic KRAS^G12D^ mutation. As observed in KRAS^G12C^ allografts, our results demonstrated that blocking SOS1-mediated signaling, both genetically and with BI-3406, resulted in significantly diminished tumor progression (**Figs. 6A-B** and **Suppl Fig. S6A**), which correlated with a reduction in cell proliferation and ERK activation in this model **(Fig. 6C** and **Suppl Figs. S5B, S6B-C)**. Moreover, SOS1 inhibition also attenuated the response in the KRAS^G12D^-related TME **(Fig. 6C** and **Suppl Fig. S6D-F)**.

**Fig. 6.**
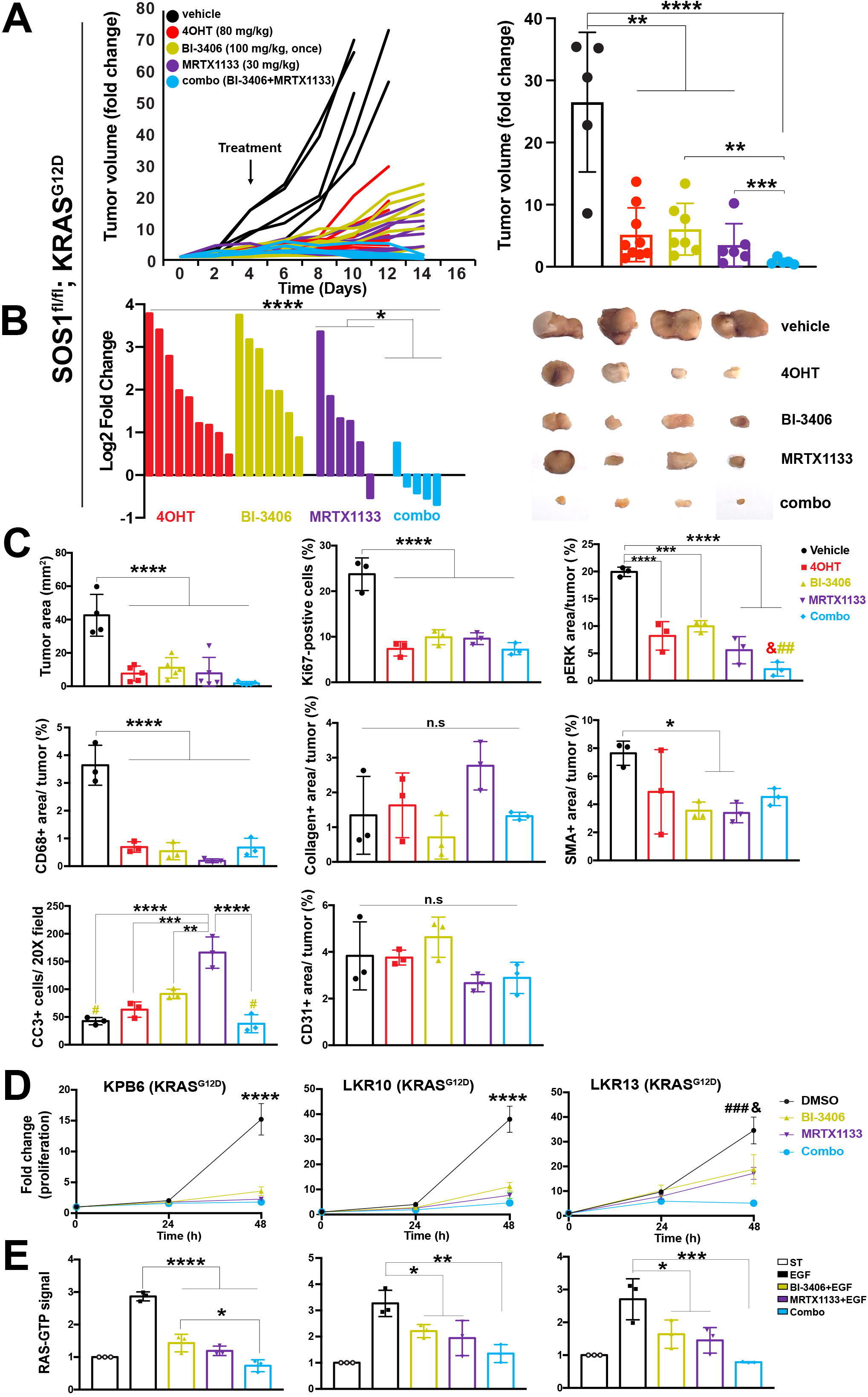
Allograft assays of (genetically or pharmacologically) SOS1-disrupted KRAS^G12D^ MEFs. **(A)** Immortalized SOS1^fl/fl^ MEFs stably expressing exogenous KRAS^G12D^ were injected subcutaneously into nude mice. Once tumors reached a size of 200 mm^3^, animals were treated (arrow) with vehicle (via gavage), 4OHT (80 mg/kg; via i.p), BI-3406 (100 mg/kg, once daily; via gavage), MRTX1133 (30mg/kg/day; i.p) or combo (BI-3406+MRTX1133; gavage/intraperitoneally) for 12 days. Data shown as mean ± SD. One-way ANOVA was used. **p < 0.01 and ****p < 0.0001. n=5 (vehicle-treated); n=9 (4OHT-treated); n=7 (BI-3406-treated); n=6 (MRTX1133-treated); n=5 (combo-treated). The mean fold-change in tumor volume relative to initial tumor volume is shown (left side). The arrow indicates treatment start. Bar graph represents tumor volume fold change relative to treatment start. *p < 0.05 and ****p < 0.0001. One-way ANOVA was used. **(B)** Waterfall plots represented as Log_2_ of the fold change of individual tumor responses from 4OHT, BI-3406, MRTX1133 or combo-treated are depicted for day 12 and day 14. Representative images of SOS1^fl/fl^/KRAS^G12D^ allografts at day 16 after tumor implantation treated (stating at day 4; arrow) with vehicle, 4OHT, BI-3406, MRTX1133 or combo (right side). *p < 0.05; ****p < 0.0001 vs vehicle-treated group. **(C)** The bar charts show total tumor area, the percentage of Ki67-positive cells, the percentage of pERK stained area with respect of total tumor area, the percentage of CD68-stained area, the percentage of collagen-stained area, the percentage of SMA-stained area, the number of CC3-positive cells per 20X field and the percentage of CD31-positive area. n=5 tumors per group (for H&E studies) and n=3 tumors per group (for the remaining assays). Data shown as mean ± SD. One-way ANOVA and Tukey’s test. **p < 0.01, ***p < 0.001 and ****p < 0.0001 vs MRTX1133-treated group; &p < 0.05 vs 4OHT-treated group; #p < 0.05 vs BI-3406-treated group. **(D)** Growth curves of KRAS^G12D^-mutated KPB6, LKR10 or LKR13 cell cultures at 24 hours and 48 hours of treatment, individually or combined, with BI-3406 (1 µM) and MRTX1133 (5 nM). DMSO was used as vehicle. n=5 independent experiments per group. Data shown as mean ± SD. One-way ANOVA and Tukey’s test. ****p < 0.0001 vs drug-treated LUAD cell lines; ###p < 0.001 vs combo-treated cells; &p < 0.05 vs MRTX1133-treated cells. **(E)** Bar chart showing the relative levels of RAS-GTP, measured by using RAS G-LISA assay, in extracts of serum-starved (ST), vehicle-pretreated, BI-3406-pretreated, MRTX1133-pretreated or combo-pretreated, for 2 hours, KPB6, LKR10 or LKR13 LUAD cells, upon EGF stimulation (100 ng/mL) for 2 min. n=3 independent samples per experimental group. Data shown as mean ± SD. *p < 0.05, **p < 0.01, ***p < 0.001, ****p < 0.0001. One-way ANOVA and Tukey’s test. CC3: cleaved caspase 3; SMA: smooth muscle actin. n.s: not significant.

### Combination therapy with BI-3406 enhances the antitumor response to a direct KRAS inhibitor

We also compared the anti-tumor effect of SOS1 inhibition with the effect of a KRAS G12D-selective inhibitor (MRTX1133) ^46^ and analyzed various parameters characterizing intrinsic and extrinsic antitumor responses in our KRAS^G12D^ allograft mouse model and in established KRAS^G12D^ cell lines.

MRTX1133 treatment (30 mg/kg) for 12 days resulted in significantly reduced tumor volume in KRAS^G12D^ allografts, which was comparable, although slightly lower, to the reduction seen with BI-3406 or 4OHT (SOS1 genetic ablation) treatments in (**Figs. 6A,B**). Of note, MRTX1133 administration had no effect on mouse body weight (**Suppl Fig. S6I**). Moreover, we also compared the pharmacokinetics (PK) profile in Hsd:Athymic Nude-Foxn1nu mice after single treatment with 50 mg/kg BI-3406 (po), single treatment with 10 mg/kg of MRTX1133 (ip), and coadministration of both compounds (Supplementary File 1). BI-3406 showed a slight reduction in area under the curve (AUC −30%) after co-administration with MRTX1133 as compared to BI-3406 alone. Exposure of MRTX1133 was unchanged compared to co-administration with BI-3406. In summary, as there was no significant alteration of the PK curve for MRTX1133 after additional treatment with or without BI-3406 a drug-drug interaction can be ruled out suggesting that the added efficacy is due to a combination effect of both compounds.

The combination of BI-3406 (100 mg/kg) with MRTX1133 (30 mg/kg) led to an even deeper reduction in tumor growth of KRAS^G12D^ allografts than observed with either treatment alone (**Figs 6A,B**), thus demonstrating a significantly enhanced anti-tumor effect resulting from the concomitant inhibition of KRAS^G12D^ and SOS1. As expected, the reduction of tumor growth caused by the single or combined drug treatments correlated with corresponding reduction of cell proliferation (Ki67) and ERK activation detected by immunohistochemistry (**Fig. 6C** and **Suppl Fig. S6B-C**) or immunoblotting (**Supp Fig. S5B**) in the treated allograft explants. Although not reaching statistical significance differences, in line with tumor growth data, single MRTX1133 administration showed reduced levels of ERK phosphorylation in comparison with 4OHT or BI-3406 treated groups (**Fig. 6C** and **Suppl Figs. S5B and S6B-C**). Importantly, a more profound impairment of tumoral RAS/MAPK signaling was always observed upon combined BI-3406 and MRTX1133 administration compared to the single drug treatments (**Fig. 6C** and **Suppl Figs. S5B and S6C**). In line with the allograft assays, single or combined BI-3406 and MRTX1133 administration in different murine KRAS^G12D^-mutated lung cancer cell lines (KPB6, LKR10 and LKR13) resulted in a potent inhibitory effect on growth of cell cultures (**Fig. 6D**) and EGF-dependent RAS activation (**Fig. 6E**).

As in SOS1-ablated (genetically or pharmacologically) KRAS^G12D^ allografts, the selective inhibition of KRAS^G12D^ by MRTX1133 also resulted in downmodulation of pro-tumorigenic TME components such as CAFs, tumor associated macrophages in comparison to vehicle-treated allografts (**Fig. 6C** and **Suppl Figs. S6D-F**).

Single MRTX1133 treatment led to noticeably increased level of apoptosis (**Fig. 6C** and **Suppl Fig. S6G**) in KRAS^G12D^ allografts at the end of study, whereas tumor samples from animals treated with MRTX1133 and BI-3406 showed similar level of apoptosis to vehicle-treated allografts (**Fig. 6C** and **Suppl Fig. S6G**).

In contrast, neither SOS1 ablation (upon 4OHT or BI-3406 administration) nor MRTX1133-mediated KRAS^G12D^ inhibition (single or combined) resulted in a modification in the percentage content of intratumor blood vessels (**Fig. 6C** and **Suppl Fig. S6H**).

In summary, BI-3406 potently synergizes with the selective KRAS^G12D^ inhibitor MRTX1133, significantly enhancing its antiproliferative effects in the tumors.

### Therapeutic efficacy of *in vivo* BI-3406 and MRTX1133 treatment in a model of KRAS^G12D^-driven LUAD in immunocompetent mice

We have recently demonstrated that genetic SOS1 ablation leads to clear inhibition of tumor development in an immunologically competent KRAS^G12D^-driven LUAD mouse model ^30^. In order to compare the anti-tumor effect of genetic versus pharmacologic SOS1 inhibition, we employed the same model here to assess the effect of single BI-3406 or MRTX1133 treatments as well as the potential synergy resulting from combined BI-3406 plus MRTX1133 dual administration (**Fig. 7** and **Supp Fig. S7**).

**Fig. 7.**
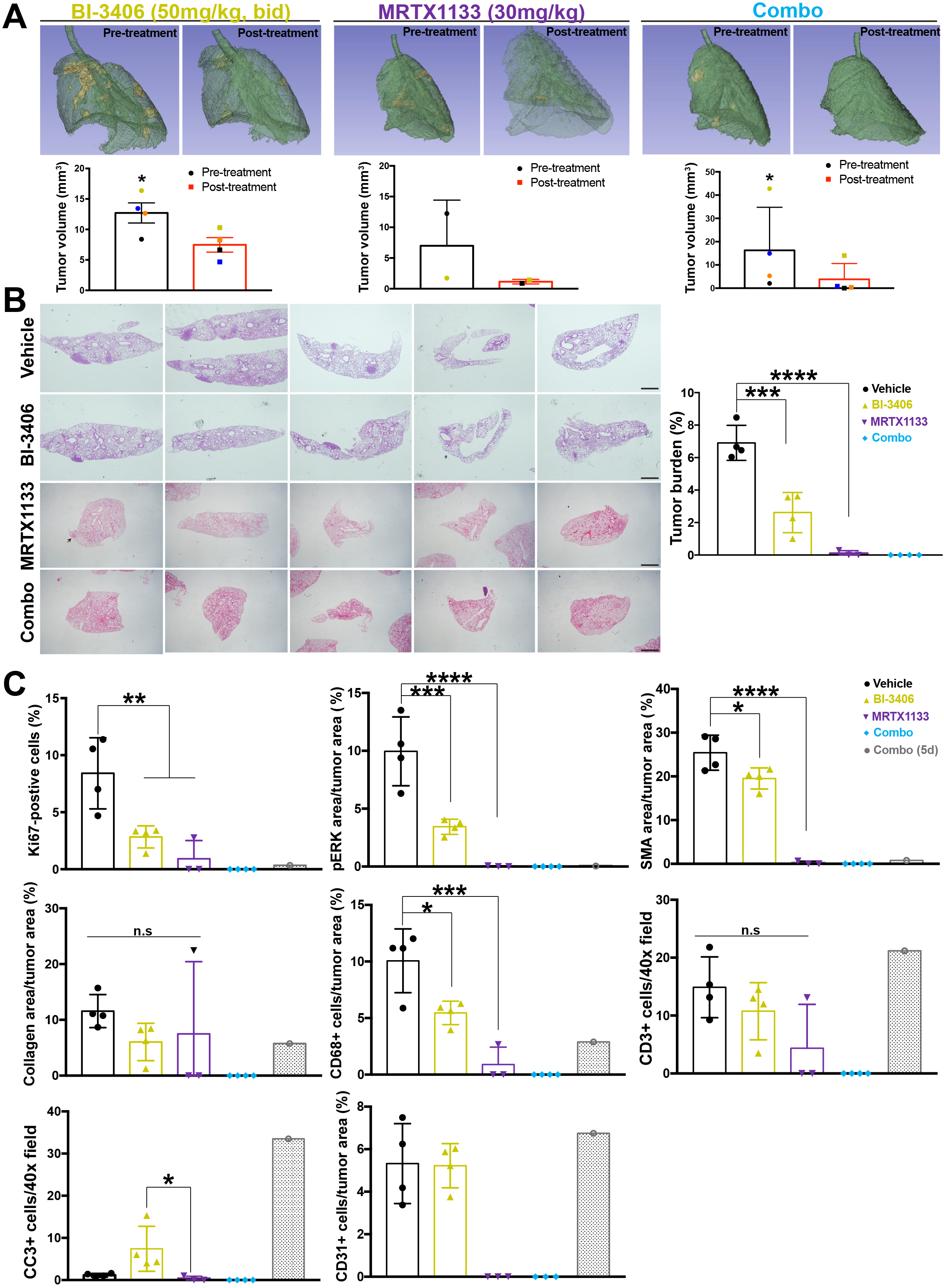
Impact of BI-3406 and MRTX1133 treatment on KRAS^G12D^-driven LUAD progression. **(A)** Representative images of microCT from 3 month-old SOS1/2^WT^/KRAS^G12D^-mutated mice previous drug treatment (designed as “pre-treatment”) and the very same animal after drug treatment (designed as “post-treatment) with BI-3406 administration (50 mg/kg, bid; via gavage, 26 days), MRTX1133-treatment (30 mg/kg, i.p, 17 days) or combo administration (BI-3406, 50mg/kg, bid; via gavage and 30mg/kg, ip for 21 days). The graphs data point corresponding to each individual mouse are identified by a distinctive color in each case. n=4 mice per genotype (BI-3406-treated); n=2 mice per genotype (MRTX1133-treated); n=4 mice per genotype (combo-3406-treated). Data shown as mean ± SD. Paired t-test was used. *p < 0.05. **(B)** Left. Representative images of paraffin-embedded sections from lung lobes of 4-month-old, SOS1/2^WT^/KRAS^G12D^-mutated mice treated with vehicle (via gavage), BI-3406 (50mg/kg; via gavage; 26 days), MRTX1133 (30mg/kg; ip; 17 days) and combo (BI-3406, 50mg/kg, bid; via gavage and 30mg/kg, ip for 21 days) stained with H&E. Scale bars: 1 mm. Right. Quantitation of percentage of lung tumor area with respect total lung area. One-way ANOVA and Tukey’s test**. (C)** The bar chart shows the percentage of Ki67-positive cells into the tumor or the percentage of pERK stained area with respect of total tumor area, the percentage of total tumor area that immunostained for SMA, the percentage of collagen in the tumor, the percentage of CD68-positive cells, the number of CD3-positive cells per 40X microscopy field in the tumor, the number of CC3-positive cells per 40X microscopy field and the percentage of CD31-stained area in vehicle-treated, BI-3406-treated, MRTX1133-treated, combo-treated samples as detailed in (B) and an additional mouse after 5 days of combo treatment. Data shown as mean ± SD. n=4/group. *p < 0.05 vs BI-3406-treated group. n=4/group. (B-C) *p < 0.05, **p < 0.01, ***p < 0.001, ****p < 0.0001. Since no tumors were found in the combo-treated group (21 days; n=4) and because only one animal was measured in the 5-days combo-treated group, both experimental groups have not been considered for the statistical analysis. Data shown as mean ± SD. One-way ANOVA and Tukey’s test.

Similar to the data obtained upon SOS1 genetic depletion ^30^, the initial analysis using *in vivo* microCT scans demonstrated a significant shrinkage of lung tumor volume after single BI-3406 or MRTX1133 administration (Post-treatment condition) in comparison to the very same animals prior to drug treatment (Pre-treatment condition). Importantly, this antitumoral effect was even higher when both drugs were concomitantly applied (**Fig. 7A**). Consistent with this, histological analysis of lungs from KRAS^G12D^ tumor-bearing animals demonstrated a reduction of lung tumor burden of more than 50% in BI-3406-treated animals as compared to vehicle-treated mice of the same age and genotype (**Fig. 7B**). Tumor reduction was stronger in MRTX1133 treated mice with only one small tumor found in the lung lobes of a total of three animals **(Fig. 7B)**. Remarkably, complete pathological response was observed in this model following co-treatment of BI-3406 and MRTX1133 as we were not able to detect remaining tumor cells in the explanted tissue **(Fig. 7B)**. Consistent with our prior observations in this animal model upon SOS1 knock-out ^30^, the lung tumor shrinkage upon BI-3406 treatment was associated with a noticeable reduction of cell proliferation (**Fig 7C** and **Supp Fig S7A**) and ERK phosphorylation (**Fig 7C** and **Supp Fig S7B**). Notice that, since mice showed complete pathological response following treatment for 26 days following co-administration with BI-3406 plus MRTX1133, we evaluated biomarker modulation in the combination following only 5 days of treatment (**Figure 7C**).

We also explored the potential effects of BI-3406 treatment on modulation of the TME of the KRAS^G12D^ lung tumors. Consistent with our previous data ^30^, immunochemical characterization of lung tumor sections derived from the BI-3406-treated mice further confirmed the positive therapeutic impact of SOS1 pharmacological inhibition with BI-3406 by identifying a clear *in vivo* downmodulation of various pro-tumorigenic components of the TME **(Fig. 7C** and **Suppl Fig. S7C-E).** We observed a modification of the morphology of the SMA-immunopositive CAFs in the tumors as a consequence of BI-3406 treatment, suggesting a BI-3406-dependent reduction of CAF activation (**Fig. 7C** and **Suppl Fig. S7C**). Furthermore, the levels of CAF-dependent collagen deposition (**Fig. 7C** and **Suppl Fig. S7D**) and CD68-reactive macrophages (**Fig. 7C** and **Suppl Fig. S7E**) present in the TME of the tumors were also strongly reduced in the lungs of BI-3406-treated as compared to vehicle-treated mice. No differences in the number of CD3-positive cells were observed upon BI-3406 single administration (**Fig. 7C** and **Suppl Fig. S7F**). MRTX1133 administered alone did not alter the number of infiltrating T-cells either (**Fig. 7C** and **Suppl Fig. S7F**), differing from other recent data carried out an oncogenic KRAS^G12D^ pancreatic cancer model showing an increase of T-cell tumor infiltration following MRTX1133 treatment ^47^. Only slightly higher (but not statistically significant) levels of CC3-positive cells were detected in the BI-3406-treated tumors as compared to vehicle-treated tumors (**Fig. 7C** and **Suppl Fig. S7G**), suggesting that the antitumoral effect of BI-3406 against KRAS^G12D^-driven lung tumors does not involve a significant priming of apoptosis in tumor cells.

Our previous results indicate that single MRTX1133 treatment increases the number of apoptotic cells, whereas the combo treatment resulted in a lower number of CC3+ cells (**Fig 6C**). Due to the enhanced efficacy observed in combination, we hypothesize that the combined MRTX1133 plus BI-3406 treatment induces apoptosis early on and therefore, at the time of evaluation at the end of the study, many apoptotic cells could have already been cleared. In this regard, our measurements of intratumor cell death rates in the experiment involving a short-term (5 days) treatment showed an increase in the number of CC3-positive cells at the initial steps of the combination treatment when compared with single MRTX1133-treated mice (**Fig. 7C** and **Suppl Fig. S7G**), thus providing first support of the above interpretation of the data in Figure 6. No differences in content of intratumor blood vessels were observed among experimental groups (**Fig. 7C** and **Suppl Fig. S7H**).

Consistent with the data previously obtained in the PDX models, these results clearly demonstrate a potent antitumoral *in vivo* effect of BI-3406, exerted through both reduction of intrinsic tumor burden and impairment of extrinsic TME in an immunocompetent KRAS^G12D^-mutated mouse model of lung adenocarcinoma.

### BI-3406 and MRTX1133 exhibit synergistically therapeutic activity in KRAS^G12D^ LUAD human cell lines

Having determined the antiproliferative effect, both *in vitro* and *in vivo*, of BI-3406 in KRAS mutant MEFs and murine cell lines and its strong synergistic effect with the mutant selective KRAS^G12D^ inhibitor MRTX1133, we next aimed at extending these observations in KRAS^G12D^ mutant human LUAD cell lines. Single BI-3406 or MRTX113 treatment did not strongly affect the proliferation and viability of A427 and SK-LU-1 human LUAD cell lines (**Fig 8A** and **Suppl Fig. S8A**). However, both compounds exhibited synergistic effects on the viability of both tumor cell lines when concomitantly administered (**Suppl Fig. S8B**). Combined BI-3406 and MRTX1133 resulted in a potent inhibitory effect on growth of those cell cultures (**Fig. 8A**). Consistently, immunoblotting analysis of human LUAD cells revealed a moderate reduction of GTP-loaded RAS as well as of ERK and Akt phosphorylation upon single BI-3406 or MRTX1133 treatment, with a more pronounced effect when both inhibitors were concomitantly administered **(Fig. 8B).**

**Fig. 8.**
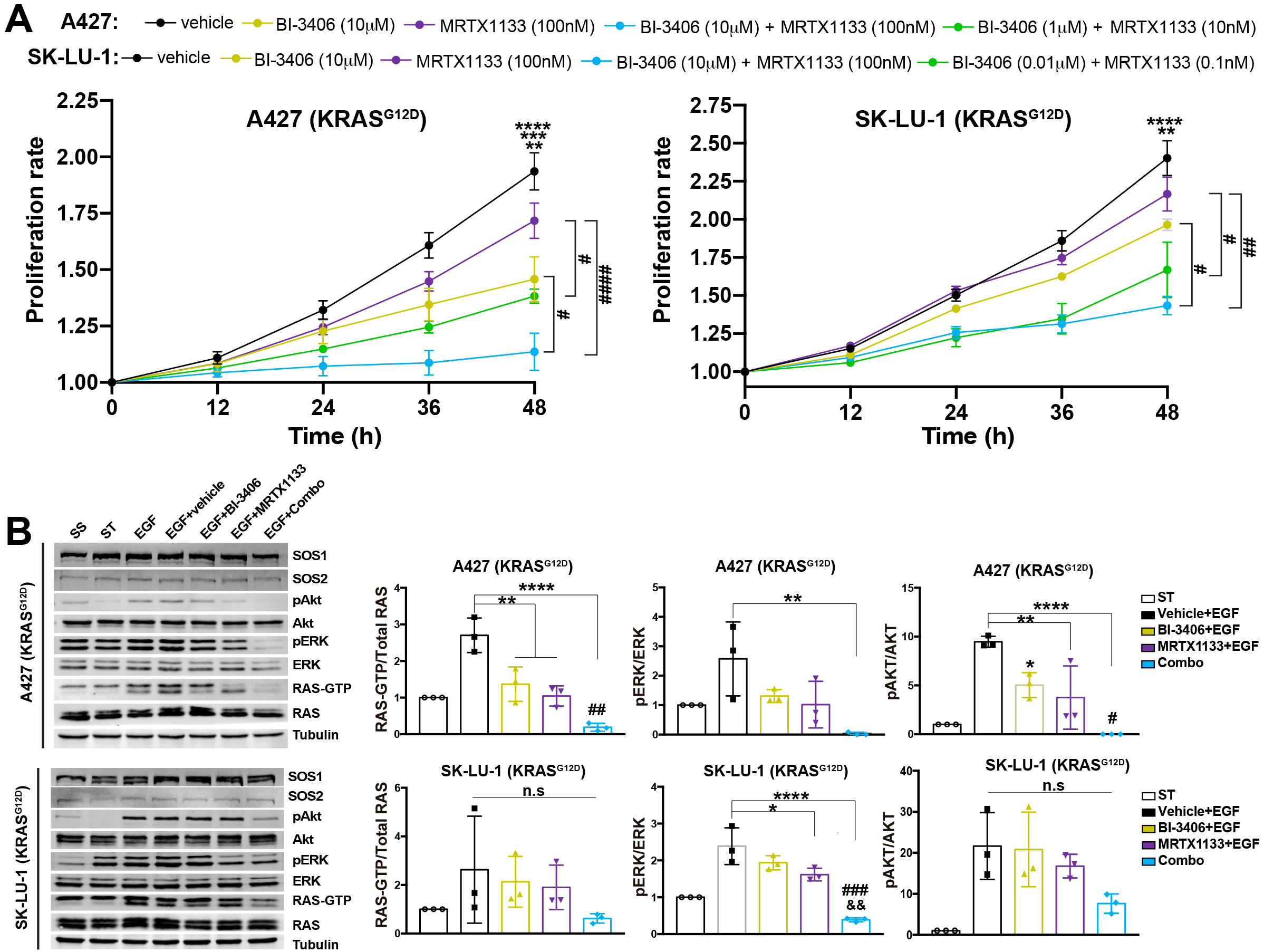
Determination of BI-3406 and MRTX113 sensitivity in KRAS^G12D^ mutant human LUAD cell lines. **(A)** Growth curves of KRAS^G12D^-mutated A427 or SK-LU-1 cells at 12, 24, 36 and 48 hours of the treatment, individually or combined, with BI-3406 and MRTX1133 at the indicated concentrations. DMSO was used as vehicle. n=5 independent experiments per group. Data shown as mean ± SEM. For A427 cells: **p < 0.01 vs BI-3406-treated (10 μM) cells; ***p < 0.001 vs BI-3406/MRTX1133-treated cells (1 μM/10 nM); ****p < 0.0001 vs BI-3406/MRTX1133-treated (10 μM/100 nM) cells. #p < 0.05 MRTX1133 (100 nM) vs BI-3406/MRTX1133-treated cells (1 μM/10 nM) or BI-3406-treated (10 μM) vs BI-3406/MRTX1133-treated (10 μM/100 nM); ####p < 0.0001 MRTX1133 (100 nM) vs BI-3406/MRTX1133-treated cells (10 μM/100 nM). For SK-LU-1: **p < 0.01 vs BI-3406/MRTX1133-treated (0.01 μM/0.1 nM) cells; ****p < 0.0001 vs BI-3406/MRTX1133-treated (10 μM/100 nM) cells; #p < 0.05 MRTX1133 (100 nM) vs BI-3406/MRTX1133-treated cells (0.01 μM/0.1 nM) or BI-3406-treated (10 μM) vs BI-3406/MRTX1133-treated (10 μM/100 nM); ##p < 0.01 MRTX1133 (100 nM) vs BI-3406/MRTX1133-treated cells (10 μM/100 nM). One-way ANOVA and Tukey’s test. **(B) (left)** Representative Western-blots of active RAS pull down assays (RAS-GTP) and the corresponding signaling of pERK and pAKT, as well as the expression of SOS1 and SOS2 in each case. Tubulin was used as a loading control. **(right)** Bar charts showing the relative levels of RAS-GTP, measured by using pull-down assay, pERK and pAkt in extracts of serum-starved (ST), vehicle-pretreated (EGF), BI-3406-pretreated, MRTX1133-pretreated or combo-pretreated (for 2 hours) A427 or SK-LU-1 cells, upon EGF stimulation (100 ng/mL) for 2 min. n=3 independent samples per group. Data shown as mean ± SD. *p < 0.05, **p < 0.01, ****p < 0.0001; &&p < 0.01 vs MRTX1133-treated cells; #p < 0.05, ###p < 0.001 vs BI-3406-treated cells. One-way ANOVA and Tukey’s test.

## DISCUSSION

Previous reports have documented the functional importance of SOS1 and SOS2 in normal cell development ^21–26^ and in the development of various tumor types ^27–31,48^ supporting the notion of SOS inhibition as a potentially useful therapeutic approach for RAS-dependent malignancies. However, the rapid lethality of genetic double-null SOS1^KO^/SOS2^KO^ mice and the observation that SOS2 can only provide limited functional compensation for the lack of SOS1 under physiological conditions, warranted further investigation of the potential impact of pharmacological SOS1 inhibition in the presence or absence of SOS2. Here, we evaluated the tolerability and therapeutic efficacy of single or combined SOS1 and SOS2 inhibition by comparing genetic or pharmacological (BI-3406 treatment) inhibition of SOS1 in SOS1/2^WT^ and SOS2^KO^ tumor cells and in genetically engineered mouse tumor models. We also compared the anti-tumor effect of genetic or pharmacological inhibition of SOS1 with a KRAS^G12D^-specific inhibitor in monotherapy and in combination with SOS1 inhibitor.

In stark contrast to the rapid lethality suffered by genetically engineered SOS1/2 double knockout mice ^21^, the inhibition of SOS1 with BI-3406 did not affect animal survival and only exhibited minimal physiological effects in SOS2^KO^ animals that were not observed in BI-3406-treated SOS1/2^WT^ mice. Furthermore, our data indicate that pharmacological SOS1 inhibition results in significant intrinsic and extrinsic therapeutic benefits by blocking the growth of the tumor cells and impairing the pro-tumorigenic effect of the TME.

An obvious mechanistic question is the reason for the observed differences between genetic and pharmacological SOS1 inhibition. BI-3406 inhibits the interaction of SOS1 with M, H, N and KRAS resulting in sensitivity of the tumor cell lines dependent on these RAS GTPases ^35,42^. In contrast, SOS1 genetic depletion (TAM-, CRISPR-, shRNA or siRNA-mediated) or SOS1 degradation removing the entire protein, not only disrupts the SOS1::RAS interaction with both oncogenic and WT RAS isoforms ^12,22,27,31,39–41,48^, but likely also affects the interaction between SOS1 and RHO GTPases, such as RAC ^28,49,50^. In this context, genetic SOS1 disruption in SOS1/2^DKO^ cells resulted in a significant decrease in RAC GTP levels ^22^, while an effect on RAC-mediated signaling after pharmacological SOS1 modulation has not yet been studied. Furthermore, BI-3406 is quickly metabolized in the organism ^35^, whereas SOS1 depletion is sustained over time.

Taken together, while genetic SOS1 depletion has stronger physiological impact in comparison to BI-3406-dependent pharmacologic SOS1 inhibition, it can lead to deleterious biological outcomes, especially in the absence of SOS2, an effect that was not detected *in vivo* upon BI-3406 administration in healthy, non-tumor bearing mice. This could be especially relevant in the context of targeted SOS1 therapies that do not just inhibit but degrade the protein, such as SOS1 PROTACs, which have recently been described ^40,41,51^. In this regard, it could be argued that systemic SOS1 inhibition may exert toxicity in humans. As previously demonstrated, SOS1 and SOS2 show some functional redundancy ^21^ and therefore the presence of SOS2 in SOS1i-treated patients could attenuate potential harmful consequence of SOS1 inhibition. In addition, our PK studies demonstrated that BI-3406 is rapidly cleared from the plasma, and therefore BI-3406-dependent SOS1 inhibition is not sustained over time, thus reducing the potential deleterious effects of a sustained SOS1 depletion.

Both in the allograft assays with KRAS^mut^ transformed MEFs and in the *in vivo* KRAS^G12D^ LUAD model in immunocompetent mice, the genetic and pharmacologic inhibition of SOS1 resulted in a significant reduction of intrinsic primary tumor growth and cell proliferation that directly correlated with decreased levels of activation of RAS/MAPK signaling pathway, in line with previous observations ^12,30,38,42,52^. Here, we found that the strong anti-tumor effect of BI-3406-mediated SOS1 inhibition was not restricted to the tumor tissue, but also impacted the surrounding stroma. In particular, we observed a significant reduction in CAF activation and in the number and collagen deposition of macrophages recruited to the tumor. Previously, BI-3406 has been shown to modulate macrophages in the TME of NF1-driven neurofibroma ^43^. These observations are also consistent with our prior data showing that genetic SOS1 depletion in LUAD mouse models reduced the activation of CAFs and tumor-associated macrophages in the TME ^30^. It should be noted that these results in LUAD differ from previous studies in models of pancreatic cancer reporting that depletion of CAFs lead to aggressive tumors and poor survival ^53,54^. Nevertheless, it has been also reported that oncogenic KRAS promotes CAFs transformations promoting tumor progression and resistance ^55–57^ and therefore, inhibiting the oncogenic signal might attenuate these pro-tumorigenic effects of CAF population. This discrepancy could reflect tissue-specific or cancer type-specific differences in responses of CAFs following SOS1/KRAS^G12D^ inhibition. Our results in LUAD also correlate with the previously characterized role of SOS1 in the physiological maintenance/homeostasis of various cell types that may be eventually recruited to the TME, including fibroblasts, macrophages, neutrophils or lymphocytes ^21–24,26,27,58^. It could be argued that that the decrease of infiltrating CAFs is due to an intrinsic effect of SOS1 inhibition, with this reduction not having a deleterious effect on tumor progression or aggressiveness in our model. Taken together, BI-3406 may elicit a dual anti-tumor effect through its antiproliferative role in the tumor cells and its potential modulation of the pro-tumorigenic TME, as observed with the selective KRAS^G12D^ inhibitor ^59^.

Interestingly, our results showed differential susceptibility of different KRAS mutants to BI-3406 treatment. In particular, BI-3406 exhibited enhanced anti-proliferative effects in KRAS^G12C^ cells, likely because this KRAS allele has a higher intrinsic GTPase activity than other KRAS mutants ^60^, and therefore is more dependent on SOS1 GEF activity to maintain the GTP-state. Interestingly, G12C cells showed incomplete SOS1 depletion even after long exposure to TAM in a SOS2^KO^ context, indicating positive selection of cells with inefficient Cre-dependent recombination. Consistent with the *in vitro* studies, SOS1 inhibition (upon TAM administration or following BI-3406 treatment) also limited tumor growth *in vivo* in allograft models expressing G12C or G12D human oncogenic variants of KRAS. In this context, BI-3406 treatment exerted stronger therapeutic effect than TAM exposure, likely due to positive selection of KRAS^mut^ cells retaining SOS1 as a result of incomplete recombination, similarly to the *in vitro* setting. Moreover, BI-3406 significantly reduced lung tumor size in a murine model of KRAS^G12D^-driven LUAD, confirming previous observations with genetically-mediated SOS1 inhibition in the same model ^30^. Similarly, recent works reported that BI-3406 also exhibited anti-tumor properties in different allograft models expressing KRAS^G12C^, demonstrating a wider therapeutic impact of targeting SOS1 in KRAS-dependent cancers ^42^.

Despite promising preclinical and clinical activity of selective KRAS^G12C^ and KRAS^G12D^ targeting agents ^61^, the emergence of drug resistance ^5–8,62^ is an important concern for most of these new small molecules ^15^. Combined administration of inhibitors that co-target upstream or downstream components of the RAS signalling pathway appears to be a promising strategy to enhance the efficacy of RAS inhibitors or to overcome resistance to these compounds ^14,42,63–65^. In fact, to address this challenge, KRAS^G12C^ inhibitors are currently tested in clinical trials in combination with drugs targeting fundamental nodes in the RAS signaling pathways, including SOS1 (NCT05578092). Genetically, we and others have demonstrated a therapeutic effect of SOS1 depletion in KRAS^G12D^-driven tumours, but the effect of combined pharmacological SOS1 and KRAS^G12D^ inhibition had not been previously evaluated. Here, we demonstrated a potent antitumor effect of the combination of BI-3406 and the selective KRAS^G12D^ inhibitor MRTX1133, which has recently entered clinical trials in advanced solid tumors (NCT05737706). Overall, our observations strongly support the consideration of SOS1 as an actionable target for pharmacological intervention in the context of RAS-dependent cancers, either as monotherapy or in combination with other drugs.

Finally, the results presented here and previously suggest that combined SOS1/SOS2 (pharmacologic/genetic) inhibition leads to anti-tumour effects in RAS-driven diseases ^12,27,30,32,48^. These observations support further evaluation of SOS2 as a potential therapeutic target for oncogenic processes *in vivo*, particularly to enhance the therapeutic effect of SOS1 inhibition. In contrast to our genetic KO model of combined SOS1/2 ablation, pharmacological inhibition of SOS1 in SOS2^KO^ mice was tolerated, suggesting that a therapeutic window may exist. Hence, the development of new drugs capable of modulating SOS1 and/or SOS2 activity *in vivo* could be a promising therapeutic avenue for the near future.

## FUNDING

Instituto de Salud Carlos III-Ministerio de Ciencia e Innovación grant FIS PI19/00934 (ES)

Junta de Castilla y León grant SA264P18-UIC 076 (ES)

Fundación Ramón Areces grant CIVP19A5942 (ES)

Instituto de Salud Carlos III -CIBERONC grant CB16/12/00352 (ES)

Asociación Española Contra el Cancer Excellence program STOP RAS CANCERS grant EPAEC222641CICS (ES)

MICIU/AEI/10.13039/501100011033/ grant PID2022-136409OB-I00 (FCB)

MICIU/AEI/10.13039/501100011033/ and European Union NextGenerationEU/PRTR grant CNS2022-135292 (FCB)

Fundación Solorzano-Barruso grant FS/32-2020 (FCB)

Fundación Eugenio Rodríguez Pascual (FCB)

Asociación Inés de Pablo Llorens-Grupo GETTHI (FCB)

Ministerio de Ciencia e Innovación grant RTI2018-099161-A-I00 (EC)

The Giovanni Armenise–Harvard Foundation (CA)

The European Research Council under the European Union’s Horizon 2020 research and innovation programme grant 101001288 (CA)

MIUR FARE grant R207ENY9KZ (CA) AIRC under IG 2021 - ID. 25737 (CA)

This research was co-financed by FEDER funds. These CIC groups are supported by the Programa de Apoyo a Planes Estratégicos de Investigación de Estructuras de Investigación de Excelencia of Castilla y León autonomous government (CLC-2017-01). MKD was supported by Fondazione Umberto Veronesi. C.A. is supported by the Zanon di Valgiurata family through Justus s.s.

The research project was selected and funded for 2 years by Boehringer Ingelheim in a Molecule for Collaboration (M4C) call submitted to the opn.me platform (www.opnme.com)

## ACKNOWLEDGMENTS

The authors thank Astrid Jeschko for her contribution performing the Pharmacokinetic studies.

## AUTHOR CONTRIBUTIONS

Conceptualization: F.C.B, C.A, E.S, B.M, K.K, M.H.H

Methodology: F.C.B, M.K.D, R.G.N, P-R.R, E.Pa, E.Pe, H.A, A.O.SJ, J.B, N.C, E.C

Investigation: F.C.B, M.K.D, R.G-N, P-R.R, E.Pa, E.Pe, H.A, A.O.SJ, E.C, C.A, E.S

Resources: J.B, E.C, B.M, K.K, M.H.H, C.A, E.S

Visualization: F.C.B, M.K.D, R.G-N, P-R.R, E.Pa, E.Pe, A.O.SJ, J.B, E.C, C.A, E.S

Funding acquisition: F.C.B, E.C, B.M, K.K, M.H.H, C.A, E.S

Project administration: C.A, E.S, B.M, K.K, M.H.H

Supervision: C.A, E.S, B.M, K.K, M.H.H

Writing – original draft: F.C.B, C.A, E.S, B.M, K.K, M.H.H

Writing – review & editing: All authors

## STAR METHODS

### KEY RESOURCES TABLE

| REAGENT or RESOURCE | SOURCE | IDENTIFIER |
| --- | --- | --- |
| Antibodies |  |  |
| Rabbit polyclonal anti-AKT | Cell Signaling Technology | Cat#9272; RRID:AB_329827 |
| Mouse monoclonal anti-AKT (40D4) | Cell Signaling Technology | Cat#2920; RRID:AB_1147620 |
| Rabbit monoclonal anti-phospho-Akt (Ser473) (D9E) | Cell Signaling Technology | Cat#4060; RRID:AB_2315049 |
| Rabbit polyclonal anti-CD3 | Abcam | Cat#ab5690; RRID:AB_305055 |
| Rabbit recombinant monoclonal anti-CD31 (EPR17259) | Abcam | Cat# ab182981; RRID:AB_2920881 |
| Rabbit polyclonal anti-CD68 | Abcam | Cat#ab125212; RRID:AB_10975465 |
| Rabbit monoclonal anti-cleaved caspase-3 (D3E9) | Cell Signaling Technology | Cat#9579; RRID:AB_10897512 |
| Mouse monoclonal anti-HA-Tag (6E2) | Cell Signaling Technology | Cat#2367; RRID:AB_10691311 |
| Rabbit polyclonal anti-HSP90 | Cell Signaling Technology | Cat#4874; RRID:AB_2121214 |
| Rabbit monoclonal anti-KI67 (SP6) | Master Diagnostica | Cat# MAD-000310QD |
| Rabbit polyclonal anti-phospho-p44/42 MAPK (Erk1/2) (Thr202/Tyr204) | Cell Signaling Technology | Cat#9101; RRID:AB_331646 |
| Mouse monoclonal anti-phospho-ERK (E-4) | Santa Cruz Biotechnology | Cat#sc-7383; RRID:AB_627545 |
| Rabbit polyclonal anti-p44/42 MAPK (Erk1/2) | Cell Signaling Technology | Cat#9102; RRID:AB_330744 |
| Mouse monoclonal anti-RAS (RAS10) | Sigma-Aldrich | Cat#05-516;<br>RRID:<br>AB_2536664 |
| Mouse monoclonal anti-SMA (1A4) | Sigma-Aldrich | Cat# A5228;<br>RRID:<br>AB_262054 |
| Rabbit polyclonal anti-SOS1 | Cell Signaling<br>Technology | Cat#5890; RRID:<br>AB_10626628 |
| Rabbit polyclonal anti-SOS2 (H-80) | Santa Cruz<br>Biotechnology | Cat#sc-15358;<br>RRID:<br>AB_2192446 |
| Rabbit polyclonal anti-phospho-S6<br>Ribosomal Protein (Ser240/244) | Cell Signaling<br>Technology | Cat#2215; RRID:<br>AB_331682 |
| Mouse monoclonal anti-beta Tubulin | Sigma-Aldrich | Cat#T5293;<br>RRID:<br>AB_477580 |
| Rabbit monoclonal anti-S6 Ribosomal<br>Protein (5G10) | Cell Signaling<br>Technology | Cat#2217; RRID:<br>AB_331355 |
| Rabbit polyclonal anti-Vinculin | ProteinTech | Cat#26520-1-AP;<br>RRID:<br>AB_2868558 |
| ECL Sheep anti-Mouse IgG HRP-linked | GE Healthcare | Cat#NA931;<br>RRID:<br>AB_772210 |
| ECL Donkey anti-rabbit IgG, HRP-linked | GE Healthcare | Cat#NA934;<br>RRID:<br>AB_772206 |
| **Bacterial and virus strains** |  |  |
| pBABE-hygro-HA-tagged | Provided by D.<br>Santamaria | N/A |
| pBABE-puro-SV40LT | Addgene | Cat#13970;<br>RRID:<br>Addgene_13970 |
| phCMV-coGALVenv-C4070A | Addgene | Cat#172217;RRID:<br>Addgene_172217 |
| pCMV-VSV-G | Addgene | Cat#8454;RRID:<br>Addgene_8454 |
| pAmpho | Ambrogio et al. <sup>10</sup> |  |
| **Chemicals, peptides, and recombinant proteins** |  |  |
| BI-3406 | Provided by<br>Boehringer-Ingelheim | N/A |
| MRTX1133 | Provided by<br>Boehringer-Ingelheim | N/A |
| Captisol | Provided by Boehringer-Ingelheim | N/A |
| Natrosol | Provided by Boehringer-Ingelheim | N/A |
| Epidermal Growth Factor | Sigma-Aldrich | Cat#E1257 |
| Tamoxifen | Harlan | Teklad CRD TAM400/CreER |
| 4-hydroxytamoxifen | Millipore | Cat#H6278 |
| Polybrene | Santa Cruz Biotechnology | Cat#sc-134220A |
| Plasmocin®- Mycoplasma Elimination Reagent | Invivogen | Ant-mpt-1 |
| Puromycin | Sigma-Aldrich | Cat#P8833 |
| AlamarBlue™ Cell Viability Reagent | Invitrogen | Cat#DAL1100 |
| CellTiter-Glo Luminescent Cell Viability Assay | Promega | Cat#G9242 |
| Effectene Transfection Reagent | Qiagen | Cat#301425 |
| JetPEI® transfection reagent | Polyplus | Cat#101000020 |
| Critical commercial assays |  |  |
| RAS G-LISA™ assay | Cytoskeleton Inc | Cat#BK131 |
| QuikChange XL Site-Directed Mutagenesis Kit | Agilent | Cat#200516 |
| Experimental models: Cell lines |  |  |
| A427 | ATCC | Cat#HTB-53 |
| SK-LU-1 | ATCC | Cat#HTB-57 |
| HEK293T | ATCC | Cat#CRL-3216 |
| KPB6 | Prof. Esther Castellano<sup>66</sup> | N/A |
| LKR10 | Prof. Esther Castellano<sup>67</sup> | N/A |
| LKR13 | Prof. Esther Castellano<sup>67</sup> | N/A |
| Mouse Embryonic Fibroblasts | Prof. Eugenio Santos<sup>22</sup> | N/A |
| Experimental models: Organisms/strains |  |  |
| *Escherichia coli* | ThermoFisher | Cat#18265017 |
| *Mus musculus*: SOS1/2 mouse colony | Prof. Eugenio Santos<sup>21</sup> | RRID:IMSR_JAX:000664 |
| *Mus musculus*: KRAS<sup>LA2</sup> mouse model | Prof. Esther Castellano<sup>67</sup> | RRID:IMSR_JAX:000664 |
| *Mus musculus*: Crl:NU(NCr)-Foxn1nu | Charles River | 490CRATHHO |
| *Mus musculus*: Hsd:Athymic Nude-Foxn1nu | Envigo RMS B.V. | Cat#6904F |
| Plasmids |  |  |
| Raf-1 GST RBD 1-149 | Addgene | Cat#13338;<br>RRID:<br>Addgene_13338 |
| Software and algorithms |  |  |
| ImageJ (v1.53t) | Schneider et al. <sup>68</sup> | https://imagej.net |
| GraphPad Prism8 | GraphPad | https://www.graphpad.com/ |
| microCT Sedecal ACQ | Baltanas et al. <sup>30</sup> | www.sedecal.com |
| Incucyte® software (v2022A) | Sartorius | https://downloads.essenbioscience.com/get/incucyte-2022a-gui |
| SynergyFinder (v3.0) | lanevski et al. <sup>69</sup> | https://synergyfinder.fimm.fi/v3/ |

### RESOURCE AVAILABILITY

#### Lead contact

Further information and requests for resources and reagents should be directed to and will be fulfilled by the lead contact, Eugenio Santos.

#### Materials availability

Further requests for cell lines generated in this study should be directed to and will be fulfilled by the lead contact, Eugenio Santos, with the Materials Transfer Agreement.

#### Data and code availability

This study did not report new original code. Any additional information required to reanalyse the data reported in this paper is available from the lead contact upon request.

### EXPERIMENTAL MODEL AND STUDY PARTICIPANT DETAILS

#### Animal models

Our conditional, tamoxifen (TAM)-inducible SOS1/2^KO^ system ^21^ involving the use of a mouse strain harboring a floxed SOS1 exon flanked by LoxP sites (SOS1^fl/fl^) controlled by Cre^ERT2^ (Cre recombinase fused to a triple mutant form of the human estrogen receptor, from the endogenous *Polr2a* locus), as well as constitutive knockout SOS2^KO^ mice ^20^, were used to generate mice of the relevant SOS genotypes (SOS1/2^WT^, SOS1^KO^, SOS2^KO^, or SOS1/2^DKO^) for these studies. All animals were kept on the same C57BL/6J background, and maintained and treated under identical experimental conditions. Genotypes were monitored by PCR as described ^21^. For SOS1 disruption in living mice, the Cre recombinase was activated by feeding the animals with TAM-containing chow diet (Harlan; Teklad CRD TAM400/CreER). For SOS1 disruption in cell cultures, the active metabolite 4-hydroxytamoxifen (4OHT) was used.

BI-3406 administration was performed as follows. For one set of animals (SOS1/2^WT^, SOS1^fl/fl^, SOS2^KO^ and SOS1^fl/fl^/SOS2^KO^ (hereafter named SOS1/2^DKO^), a TAM-containing chow diet (Harlan; Teklad CRD TAM400/CreER) was administered to 5-week-old mice (no gender selected) to achieve SOS1 depletion in SOS1^KO^ and SOS1/2^DKO^ groups. All experimental groups were TAM-fed to avoid undesirable off-target effects. Another set of 5 weeks-old mice with SOS1/2^WT^ and SOS2^KO^ genotypes were orally treated (gavage) on a 5 days on/2 days off bid (6 hours apart) schedule with vehicle (0.5% Natrosol) or BI-3406 at 50 mg/kg for 26 days as described ^35^.

For additional *in vivo* tests, SOS1/2^WT^ mice were cross-mated with KRAS^LA2^ mice spontaneously developing KRAS^G12D^-driven LUAD ^30,67^. All mice were kept on the same C57BL/6J background (RRID:IMSR_JAX:000664) and maintained under identical experimental conditions. 3 month-old mice (no gender selected) were orally treated (gavage) on a 5 days on/2 days off bid (6 hours apart) schedule with vehicle (0.5% Natrosol) and BI-3406 at 50 mg/kg for 26 days or with MRTX1133 (30 mg/kg, administered i.p.) for 17 days. Combo (BI-3406 plus MRTX1133) drug treatment was also performed for 21 days using the same treatment regimen as described for single administration.

For allograft experiments, Crl:NU-Foxn1^nu^ mice (females, 8-week-old) were purchased from Charles River. KRAS^G12C^;SOS1^fl/fl^;SOS2^+/+^;Cre^+^ or KRAS^G12D^;SOS1^fl/fl^;SOS2^+/+^;Cre^+^ cells (2×10^6^) were injected subcutaneously in the flanks of recipient mice. Once tumors were detectable, mice were randomly assigned to either BI-3406, 4OHT or vehicle treatment [For KRAS^G12C:^ n=5 (vehicle-treated); n=9 (4OHT-treated); n=10 (BI-3406-treated) and For KRAS^G12D^: n=5 (vehicle-treated); n=9 (4OHT-treated); n=10 (BI-3406-treated); n=6 (MRTX1133-treated); n=5 (combo-treated)] and measurements were taken every two days using calipers. Body weight was assessed every other day in control mice treated with MRTX1133. BI-3406 (provided by Boehringer-Ingelheim) was resuspended in 0.5% Natrosol (provided by Boehringer-Ingelheim), 1 equimolar HCl (1M) and administered daily by oral gavage at a dose of 100 mg/kg (once daily). TAM was dissolved in corn oil and administered as intraperitoneal (i.p.) injection (100 μl total volume) 4 consecutive days/week at a dose of 80 mg/kg. MRTX1133 (provided by Boehringer-Ingelheim) was dissolved in 10% Captisol in 50 mM citrate buffer pH5.0 and administered as intraperitoneal (i.p.) injection every day at a dose of 30 mg/kg. Mice were euthanized and tumors were resected. Half of the tumors were snap frozen and the second part was fixed in formalin and embedded in paraffin for further analysis.

Mice were kept, managed, and sacrificed in the NUCLEUS animal facility of the University of Salamanca or at the MBC Animal Facility of the University of Torino, according to current European (2007/526/CE) and Spanish (RD 1201/2005 and RD53/2013) legislation in accordance with the guideline for Ethical Conduct in the Care and Use of Animals as stated in The International Guiding Principles for Biomedical Research Involving Animal. The mice were housed in cages with adequate space, bedding material for comfort and maintained under specific pathogen free conditions, while maintaining 12-hour dark/light cycle. Ambient temperature was kept within 20-24°C, and humidity levels ranged from 45-65%. All experiments were approved by the Bioethics Committee of the Cancer Research Center (#596) and by the Italian Health Minister (authorization n° 1227/2020-PR).

#### Cell lines

Human lung adenocarcinoma cell lines SK-LU-1 (ATCC, Cat#HTB-57) and A427 (ATCC, Cat#HTB-53) holding KRAS^G12D^ mutation were grown in minimum essential medium (MEM) containing 10% FBS, 1% penicillin/streptomycin, 0.5 mg/mL Fungizone (Amphotericin B) and 50 µg/mL of Plasmocin® (Invivogen) at 37°C and 5% CO_2_. Murine cells were grown in Dulbecco’s modified Eagle’s medium (DMEM) (4.5 g/L glucose + 10% FBS + 1% penicillin/streptomycin) at 37°C and 5% CO_2._. The murine KPB6 cells (harboring KRAS^G12D^ and p53 mutations) were generated in Sergio Quezadás Laboratory (University College London), used by Julian Downward’s lab ^66^ and kindly provided by Dr. Esther Castellano. The LKR10 and LKR13 are mouse KRAS^G12D^-mutant lung cancer cell lines derived by serial passage of minced lung adenocarcinoma tissues from two tumors isolated from separate lobes of the Kras^LA1^ mouse model ^67^ and kindly provided by Dr. Esther Castellano. To evaluate cell proliferation, AlamarBlue™ Cell Viability Reagent (Invitrogen, Cat#DAL1100) was added (10 µL/well) and incubated in dark for 1 hour at 37°C and 5% CO_2_. Fluorescence intensity was then evaluated at 590 nm using TECAN Infinite® 200 PRO microplate reader taking measurements at 0, 24 and 48 hours. In particular, 1×10^6^ of murine KRAS^G12D^-mutant LUAD cell lines (LKR10, LKR13 or KPB6) cells were treated with vehicle (DMSO), BI-3406 (1 µM) and/or MRTX1133 (5 nM) and cell proliferation quantitated as described above. In addition, the effect of BI-3406 on RAS activation was measured using the RAS G-LISA™ assay (Cytoskeleton Inc, Cat#BK131) in serum-starved KPB6, LKR10 and LKR13 (KRAS^G12D^-mutated) LUAD cells treated with vehicle (DMSO), BI-3406 (1 µM) and/or MRTX1133 (5 nM) for 2 hours and stimulated with EGF (100 ng/mL) for 2 minutes.

#### Mouse Embryonic Fibroblast immortalization

Mouse Embryonic Fibroblasts were immortalized by introducing the SV40LT antigen in primary MEFs isolated from mice of the four SOS1/2 relevant genotypes. HEK293T cells (ATCC, Cat#CRL-3216) grown in DMEM medium (4.5g/L glucose, Gibco + 10% FBS + 1% P/S) were used to produce the retroviral vector containing the SV40LT-Antigen. 3×10^6^ HEK293T cells were seeded the day before transfection. Before transfecting HEK293T cells, the culture medium was changed to antibiotics-free DMEM. The Retroviral vector assembly formulation consisted of: 1mL Opti-MEM medium (Gibco, Cat#31985070), pBABE-puro-SV40LT (12.5 µg, Addgene, Cat#13970), phCMV-coGALVenv-C4070A (9.5 µg, Addgene, Cat#172217) pCMV-VSV-G (3 µg, Addgene, Cat#8454) and JetPEI® transfection reagent (62.5µL, Polyplus, Cat#101000020) per each p100 plate. Described mix was incubated for 20 minutes before gently adding it to the culture plate. After 48 hours, the supernatant was collected, centrifuged (5 minutes, 1500 rpm, at room temperature), and filtered (0.45 µm pore size) to obtain filtered virus.

SOS1/2^WT^, SOS1^fl/fl^, SOS2^KO^, and SOS1^fl/fl^/SOS2^KO^ primary MEFs were trypsinized and seeded at a density of 1×10^5^ cells. MEFs were then infected with 2mL of the filtered virus with Polybrene® (8µg/mL, Santa Cruz, Cat#sc-134220A) and centrifuged at 800g for 30 minutes at RT. Reinfection was carried out 24 hours later following the same methodology. 24 hours after reinfection, Puromycin (2µg/mL, Sigma, Cat#P8833) was supplemented to the culture media for positive selection of infected MEFs until complete immortalization was achieved.

#### Generation of SOS-less MEFs expressing human WT or mutant KRAS

Immortalized SOS-less MEFs including control SOS1/2^WT^ (*SOS1*^+/+^;*SOS2*^+/+^;Cre^+^), single SOS1^KO^ (*SOS1*^fl/fl^;*SOS2*^+/+^;Cre^+^), single SOS2^KO^ (*SOS1*^+/+^;*SOS2*^−/−^;Cre^+^), and SOS1/2^DKO^ (*SOS1*^fl/fl^;*SOS2*^−/−^;Cre^+^) were established as described ^21^.

KRAS^G12C^, KRAS^G12D^ and KRAS^G12V^ retroviral plasmids were created by point mutagenesis into pBABE-hygro-HA-tagged KRAS^WT^ plasmid (provided by David Santamaria) using QuikChange XL Site-Directed Mutagenesis Kit (Agilent, Cat#200516). Retroviruses were generated by co-transfecting pBABE plasmids together with pAmpho plasmid into 293T cells using Effectene Transfection Reagent (Qiagen, Cat#301425). The retroviruses were transduced into *SOS*-less MEFs followed by 2 weeks of hygromycin selection (200 μg/mL) in complete DMEM medium. To induce *SOS1* ablation, cells were then cultured for at least 11 days in the presence of 4-hydroxytamoxifen (4OHT; Sigma, 600 nmol/L, Cat#H6278). Cells were grown at 37°C and 5% CO_2_ in a humidified incubator in DMEM medium supplemented with 10% fetal bovine serum, 100 mg/mL penicillin and 100 units/mL streptomycin.

### METHOD DETAILS

#### Pharmacokinetic study

Three female Hsd:Athymic Nude-Foxn1nu mice, were used per treatment group. Mice were treated once with either 50 mg/kg BI-3406 (p.o.), 10 mg/kg MRTX1133 (i.p.) or with both compounds. Blood samples were collected from each mouse by penetrating the *vena saphena* with a needle in a tube containing EDTA as anticoagulant (Microvette CB 300 K2E, Sarstedt) at 5 timepoints within 24 hours. Mice were restrained during blood sampling. Blood was centrifugated to separate 10 µl plasma for measuring compound concentrations. Pharmacokinetic mouse study performed at Boehringer-Ingelheim were approved by the internal ethics committee and the local Austrian governmental committee. Mice used in Boehringer-Ingelheim studies are group housed within environmentally controlled conditions with a 12-hour-light/dark cycle at 21°C ± 1.5 °C, 55% ± 10% humidity, and receive food and water ad libitum.

Compound concentrations in plasma aliquots were measured by quantitative HPLC-MS/MS using an internal standard. Calibration and quality control samples were prepared using blank plasma from untreated animals. Samples were precipitated with acetonitrile and injected into a HPLC system (Agilent 1200). Separation was performed by gradients of 5 mmol/L ammonium acetate pH 4.0 and acetonitrile with 0.1% formic acid on a 2.1 mm by 50 mm Xbridge BEH C18 reversed-phase column with 2.5 µm particles (Waters). The HPLC was interfaced by ESI operated in positive ionization mode to a triple quadrupole mass spectrometer (5000 or 6500+ Triple Quad System, SCIEX) operated in multiple reaction monitoring mode. Chromatograms were analyzed with Analyst (SCIEX) and pharmacokinetic parameters were calculated by non-compartmental analysis using BI-proprietary software.

#### Histology and immunostaining

For histological examination, mice were euthanized and their organs of interest dissected and fixed in 4% of paraformaldehyde for 24 hours. Tissue blocks were then dehydrated and paraffin-embedded. Paraffin-embedded lung samples were cut (3 μm thick) and stained with hematoxylin and eosin or Masson’s trichrome according to standard procedures. For immunohistochemistry, sections were dewaxed, microwaved in citrate buffer (pH 6) and incubated overnight with anti-pERK1/2 (Cell Signaling, 1:500, Cat#9101), anti-Ki67 (SP6) (Master Diagnostica, 1:50, Cat#MAD-000310QD), anti-SMA (1A4) (Sigma, 1:500, Cat#A5228), anti-CD3 (Abcam, 1:50, Cat#ab5690), anti-CD31 (Abcam, 1:1000, Cat#ab182981), anti-CD68 (Abcam, 1:100, Cat#ab125212) and anti-CC3 (D3E9) (Cell Signaling, Cat#9579, 1:250) at 4°C. Sections were then incubated with biotin-conjugated secondary antibodies followed by Vectastain Elite ABC reagent and the reaction product visualized by incubating the sections in 0.025% 3.3’-diaminobenzidine and 0.003% H_2_O_2_ in PBS. Processing and staining of the sections, as well as the histopathological evaluation of the samples ^70^ were performed by the PMC-BEOCyL Unit (Comparative Molecular Pathology-Biobank Network of Oncological Diseases of Castilla y León).

#### Hematological and biochemical parameters

To monitor hematological parameters, blood samples were taken from the submandibular sinus at the indicated time points into microvette ethylenediaminetetraacetic acid (EDTA)-coated tubules (Sarstedt Inc., Nümbrecht, Germany). The Hemavet 950 instrument (Drew Scientific, Dallas, USA) was used to determine hematological cell counts and other parameters in peripheral blood (PB) samples obtained from the two defined experimental groups (TAM-treated or BI-3406/vehicle-treated). Various biochemical parameters in blood serum from different experimental groups of animals treated with TAM for 14 days, or BI-3406/vehicle for 26 days, were also determined using reagent strips (total bilirubin, Panel-2, Panel-V, and Heart-2 test strips; Menarini Diagnostics, Barcelona, Spain) quantified on the Spotchem EZ 4430 instrument (Menarini Diagnostics).

#### Micro CT scanning

To evaluate the impact of BI-3406 and MRTX1133 administration (individually or combined) on tumor regression, BI-3406-untreated, 3-month-old, SOS1/2^WT^/KRAS^G12D^ mice (Pre-treatment condition), and the very same mice (Post-treatment condition) just at the end of single BI-3406 treatment (50 mg/kg, bid, via gavage, 26 days), single MRTX1133-treatment (30 mg/kg, i.p, 17 days) or combo administration (BI-3406, 50mg/kg bid, via gavage and 30mg/kg, ip for 21 days) were deeply anaesthetized and imaged using the SuperArgus Micro-CT (SEDECAL, Madrid, Spain). Images were taken with 720 plane projections, 100 ms exposure time per projection and X-ray energies of 45 kVp and 400 µA. Images were reconstructed and converted to 3D volumes using microCT Sedecal ACQ software and tumoral and non-tumoral segmentations were performed by using the 3D Slicer image computing platform. Images were taken at the NUCLEUS Molecular Imaging Laboratory of the University of Salamanca.

#### IncuCyte Growth Assays

Growth rate of KRAS^G12C^, KRAS^G12D^ and KRAS^G12V^ SOS-less immortalized MEFs cells was assessed as previously described ^10^. Cells (1 × 10^3^) were seeded in 96-well plates in 100 μL DMEM complete medium. After overnight incubation, 100 μL DMEM complete medium containing BI-3406 was added to achieve a concentration of 1 μM BI-3406. The plates were then incubated in the IncuCyte S3 for real-time imaging, with four fields imaged per well under 10x magnification every two hours for a total of 86 to 90 hours. Data were analyzed using the IncuCyte 2022A software, which quantified cell surface area coverage as confluence values. IncuCyte experiments were performed in triplicate. A single representative growth curve is shown for each condition. The data were graphically displayed using GraphPad Prism 8 for Windows (GraphPad Software).

#### Drug-response assay

Cells (1 × 10^3^) were seeded in 96-well plates. The following day, cells were treated with BI-3406 using a nine-point 2-fold dilution series. After 72 hours of incubation the cell viability was measured using CellTiter-Glo Luminescent Cell Viability Assay (Promega, Cat#G9242). All experimental points were a result of three replicates. Each point (mean ± Standard deviation) represents growth of treated cells normalized to cells treated with DMSO. Inhibitory concentration values were calculated using four parameter logistic curve fitting using GraphPad Prism 8 for Windows (GraphPad Software). All experiments were repeated at least three times.

#### RAS activation pull down assay and Western-blot

SK-LU-1 or A427 cultured human cells (10^6^) were serum-starved for 24 hours and then treated with BI-3406 (1 μM), MRTX1133 (5 nM) or combo (1 μM+5 nM) for 2 hours. Vehicle-treated (0.1% v/v of DMSO) cells were used as control. The cells were then stimulated with the Epidermal Growth Factor (EGF, 100 ng/mL, Sigma-Aldrich, Cat#E1257) for 2 minutes. Similarly, immortalized MEFs of the four defined SOS1/2 genotypes expressing different KRAS mutations (G12C, G12D, G12V) were maintained in DMEM with 10% FBS and glutamine. To evaluate the effect of genetic SOS1/2 ablation, MEFs from the four experimental groups were treated with 4OHT (0.3 µM) under identical conditions to induce SOS1 removal and also exclude any possible off-target effects. 5×10^5^ cells were serum-starved by 12 hours and then stimulated with EGF (100 ng/mL) for 2 minutes. Similarly, the effect of BI-3406 on RAS activation was also evaluated in MEFs of the same experimental groups above mentioned. Cells were pre-treated with vehicle (DMSO) or BI-3406 (1 μM) for 2 hours and stimulated with EGF (100 ng/mL) for 2 minutes. Cells were scraped in ice-cold MLB (Mg^2+^ Lysis/Wash Buffer) and the lysates were centrifuged at 14.000 rpm for 10 minutes at 4°C. A 0.5-mL aliquot of each cell extract was added to 10 µg of a bacterially-expressed GST fusion protein of Raf-1 RBD (RAS binding domain, residues 1 to 149) and incubated with Glutathione-SepharoseTM 4 Fast Flow beads (Cytiva, Cat#17513201) for 1 hour at 4°C under gentle rotation. After extensive washes in lysis buffer, the protein complexes were released by boiling in SDS-PAGE sample buffer, separated electrophoretically, transferred onto PVDF membrane, and analyzed by immunoblotting using anti-panRAS antibody (Millipore, 1:1000, Cat#005-516). Aliquots of the cell lysates were used in parallel immunoblot analysis to detect the expression of the total proteins. RAS-GTP levels were quantified using as normalizing control the total levels of RAS found in each cell lysate. Membranes were imaged using the Li-Cor Odyssey Imaging System V3.0, and quantification was performed using ImageJ software (v1.53t).

Cells from *in vitro* culture or *ex vivo* explants were lysed in RIPA lysis buffer (Thermo Fisher, Cat#89900) supplemented with Halt protease and phosphatase inhibitor cocktail (Thermo Fisher, Cat#78445). 15 μg of protein extracts were separated by SDS-PAGE (Bio-Rad), transferred to a PVDF membrane, and blotted with primary antibodies raised against RAS (Cell Signaling, Cat#8832), HA-Tag (6E2) (Cell Signaling, 1:1000, Cat#2367), HSP90 (Cell Signaling, 1:1000, Cat#4874), Phospho-AKT (Ser473) (D9E) (Cell Signaling, 1:2000, Cat#4060), Phospho-p44/42 MAPK (pErk1/2) (Thr202/Tyr204) (Cell Signaling, 1:1000, Cat#9101), Phospho-ERK (Santa Cruz, 1:500, Cat#sc-7383), Phospho-S6 Ribosomal Protein (Ser240/244) (Cell Signaling, 1:1000, Cat#2215), AKT (Cell Signaling, 1:1000, Cat#9272 and Cat#2920), p44/42 MAPK (Erk1/2) (Cell Signaling, 1:1000, Cat#9102), S6 Ribosomal Protein (5G10) (Cell Signaling, 1:1000, Cat#2217), SOS1 (Cell Signaling, 1:1000, Cat#5890), SOS2 (H-80) (Santa Cruz Biotech, 1:500, Cat#sc-15358), Tubulin (Sigma-Aldrich, 1:10000, Cat#T5293) and Vinculin (ProteinTech, 1:1000, Cat#26520-1-AP). Secondary anti-mouse or anti-rabbit antibodies include ECL Sheep anti-Mouse IgG HRP-linked secondary antibody (GE Healthcare, Cat#NA931V) and ECL Donkey anti-rabbit IgG, HRP-linked secondary antibody (GE Healthcare, Cat#NA934V). Western-blots shown are representative of at least three independent experiments.

### QUANTIFICATION AND STATISTICAL ANALYSIS

GraphPad Prism 8.0.1 (GraphPad Inc., USA) software was used. Statistical significance was determined by one-way ANOVA using the Tukey’s method to correct for multiple comparisons. For comparisons established only between two groups we used Student’s two-tailed, unpaired t-test. To compare tumor growth inhibition in the very same animals, paired t-test was performed. Survival analysis was performed by the Kaplan–Meier method and between-group differences in survival were tested using the Log-rank (Mantel-Cox) test. IncuCyte experiments and Western-blots assays were performed in triplicate. *n* values mentioned in the figure legends indicate the number of animals used per experimental group. Results are expressed as mean ± Standard Deviation (SD). Significant differences are considered at *P* value < 0.05. The investigators were blinded during evaluation of tumor size variations following treatments. For dataset analysis, the natural logarithm of the half-maximal inhibitory concentration, Ln(IC50), represents drug sensitivity. SynergyFinder web application (v 3.0) ^69^ was used to calculate and represent the potential synergy of BI-3406 and MRTX1133. Synergy scores were calculated based on the Zero Interaction Potency (ZIP) model ^71^ and represented as a 2D synergy map.

