## Supplemental Figures 1 to 8 and Supplemental Tables 1 and 2 for "Pharmacological SOS1 inhibitor BI-3406 demonstrates *in vivo* anti-tumor activity comparable to SOS1 genetic ablation in KRAS mutant tumors"

**Supplementary Figure 1**

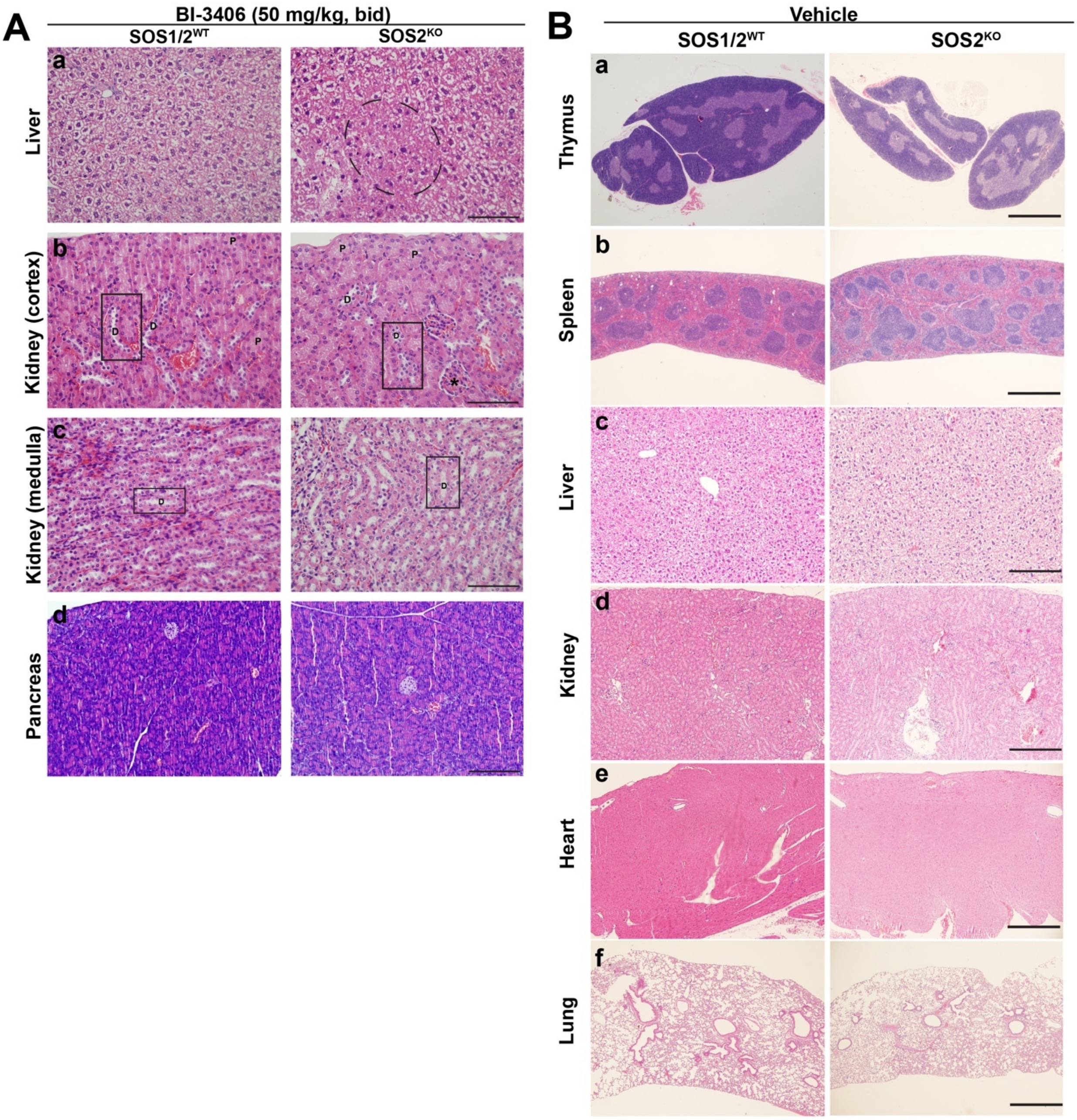


**Supplementary Fig. S1. Detailed histological examination of the impact of BI-3406-mediated SOS1 inhibition in naïve and SOS2$^{KO}$ mice. (A)** Representative images of paraffin-embedded sections from liver **(a)**, kidney (cortex and medulla; **b** and **c**, respectively) and pancreas **(d)** of BI-3406-treated (50 mg/kg, bid; via gavage; 26 days) SOS1/2$^{WT}$ and SOS2$^{KO}$ mice stained with H&E. Dashed ellipse in A (right panel) shows clusters of hepatocytes with retracted cytoplasm. Boxes indicate distal tubules. Scale bars: (a-c): 50 µm; (d): 100 µm. D and P characters in panels b and c mean Distal and Proximal, respectively. Asterisk indicates a glomerulus. **(B)** Representative images of paraffin-embedded sections stained with H&E from thymus **(a)**, spleen

**(b)**, liver **(c)**, kidney **(d)**, heart **(e)** and lung **(f)** of vehicle-treated (for 26 days) SOS1/2$^{WT}$ and SOS2$^{KO}$ mice. All treatments started 1 month of age. Scale bars: (a, b, f): 500 μm; (c-d): 100 μm; (e): 200 μm.

#### Supplementary Figure 2

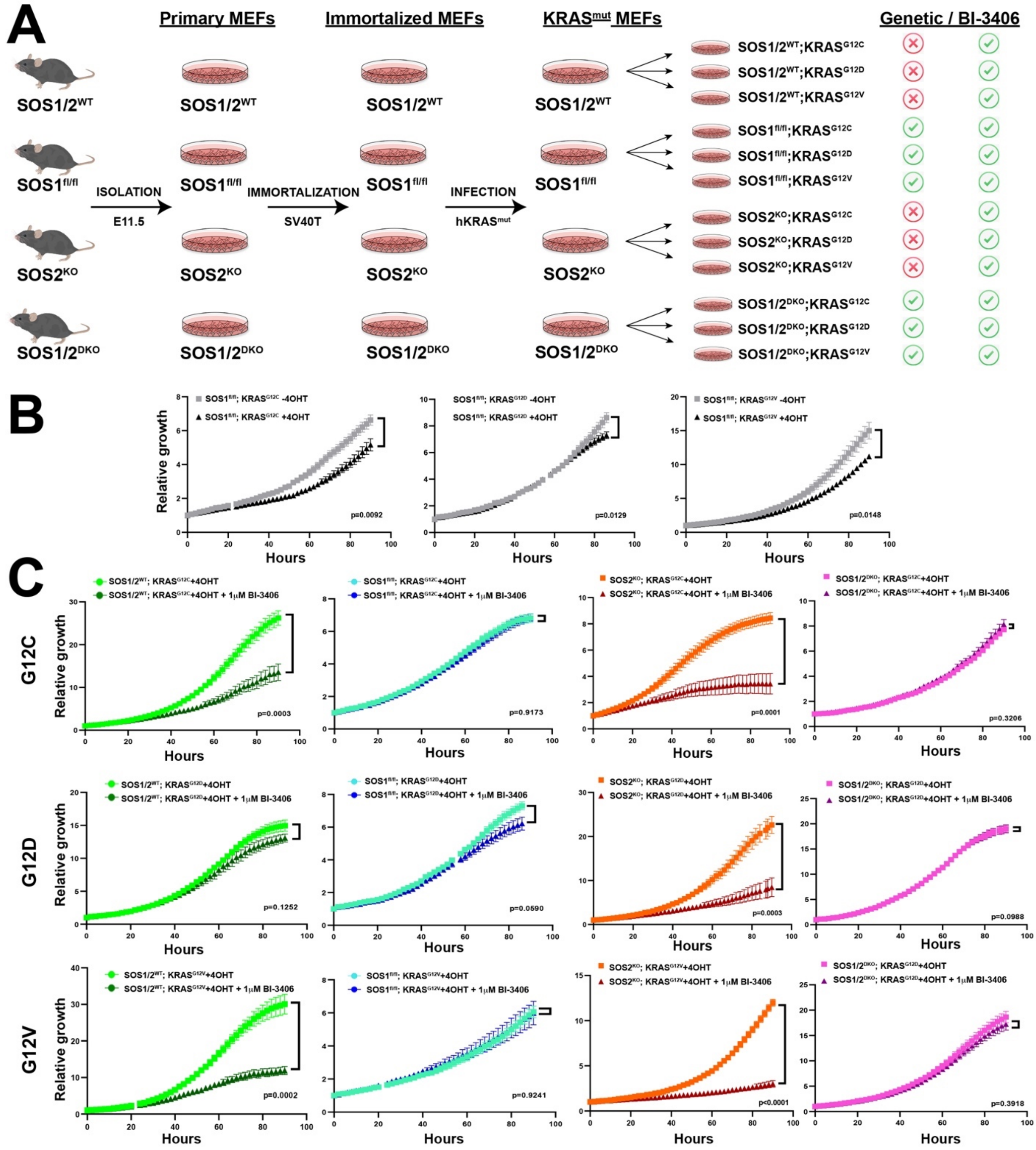


**Supplementary Fig. S2. Evaluation of genetic/pharmacologic SOS1 removal in cell proliferation and RAS/MAPK signal transmission in oncogenic KRAS mutant fibroblasts.** **(A)** Schematic illustration of the experimental design to generate oncogenic $KRAS^{mut}$ immortalized MEFs of the different SOS1/2 genotypes. **(B)** Growth curves of untreated and 4OHT-treated $SOS1^{fl/fl}$ immortalized MEFs expressing $KRAS^{G12C}$, $KRAS^{G12D}$ or $KRAS^{G12V}$ mutations as assessed by live-cell microscopy. Data expressed as mean ± SD. n=3 independent experiments per experimental group. **(C)** Growth curves of 4OHT-treated, BI-3406-treated (1 μM) or untreated, immortalized MEFs of the four SOS1/2 genotypes (SOS1/2$^{WT}$, SOS1$^{KO}$, SOS2$^{KO}$ and SOS1/2$^{DKO}$) expressing KRAS$^{G12C}$ (upper row), KRAS$^{G12D}$ (middle row) or KRAS$^{G12V}$ (lower row) as assessed by live-cell microscopy. Data expressed as mean ± SD. n=3 independent experiments per experimental group.

#### Supplementary Figure 3

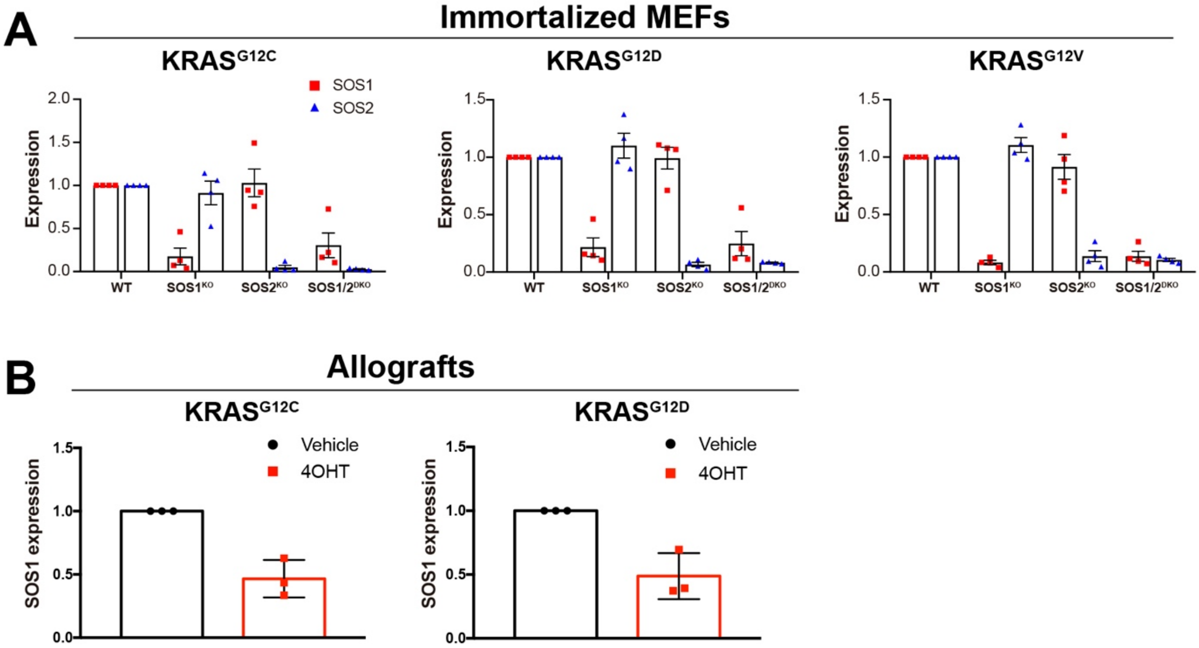


**Supplementary Figure S3**. **SOS1 protein levels in 4OHT-treated MEFs and allografts. (A)** The bar charts show SOS1 and SOS2 protein expression levels in 4OHT-treated $KRAS^{G12C}$, $KRAS^{G12D}$ and $KRAS^{G12V}$ immortalized MEFs. n=4 independent experiments. Data shown as mean ± SD. **(B)** The bar charts show SOS1 protein expression levels in isolated allografts from $SOS1^{fl/fl}/KRAS^{G12C}$ and $SOS1^{fl/fl}/KRAS^{G12D}$ mice. n=3. Data shown as mean ± SD

**Supplementary Figure 4**

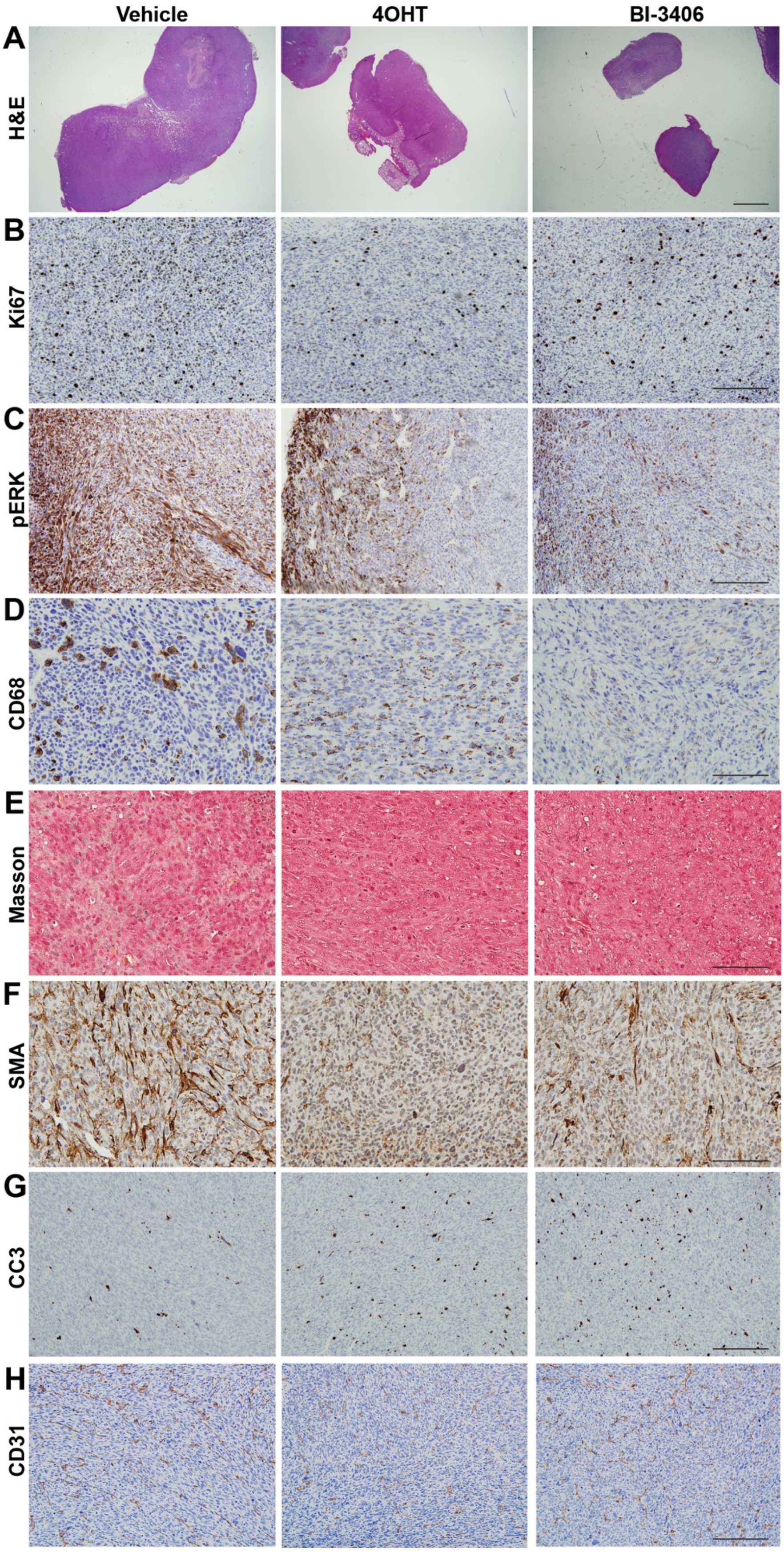

**Supplementary Figure S4. Impact of genetic/pharmacological SOS1 ablation on markers for cell proliferation, cell death and tumor microenvironment in KRAS$^{G12C}$-mutated allografts.** Representative images of paraffin-embedded sections of KRAS$^{G12C}$ mutated allografts removed from nude mice treated with vehicle, 4OHT (80 mg/kg) or BI-3406 (100 mg/kg, once daily) and stained for H&E **(A)** and Masson's trichrome **(E)** or immunostained for Ki67 **(B)**, pERK **(C)**, CD68 **(D)**, SMA **(F),** CC3 **(G)** or CD31 **(H)**. Scale bars: (A): 1 mm; (B, C, G, H): 100 μm; (D-F): 50 μm. CC3: cleaved caspase 3; SMA: smooth muscle actin.

**Supplementary Figure 5**

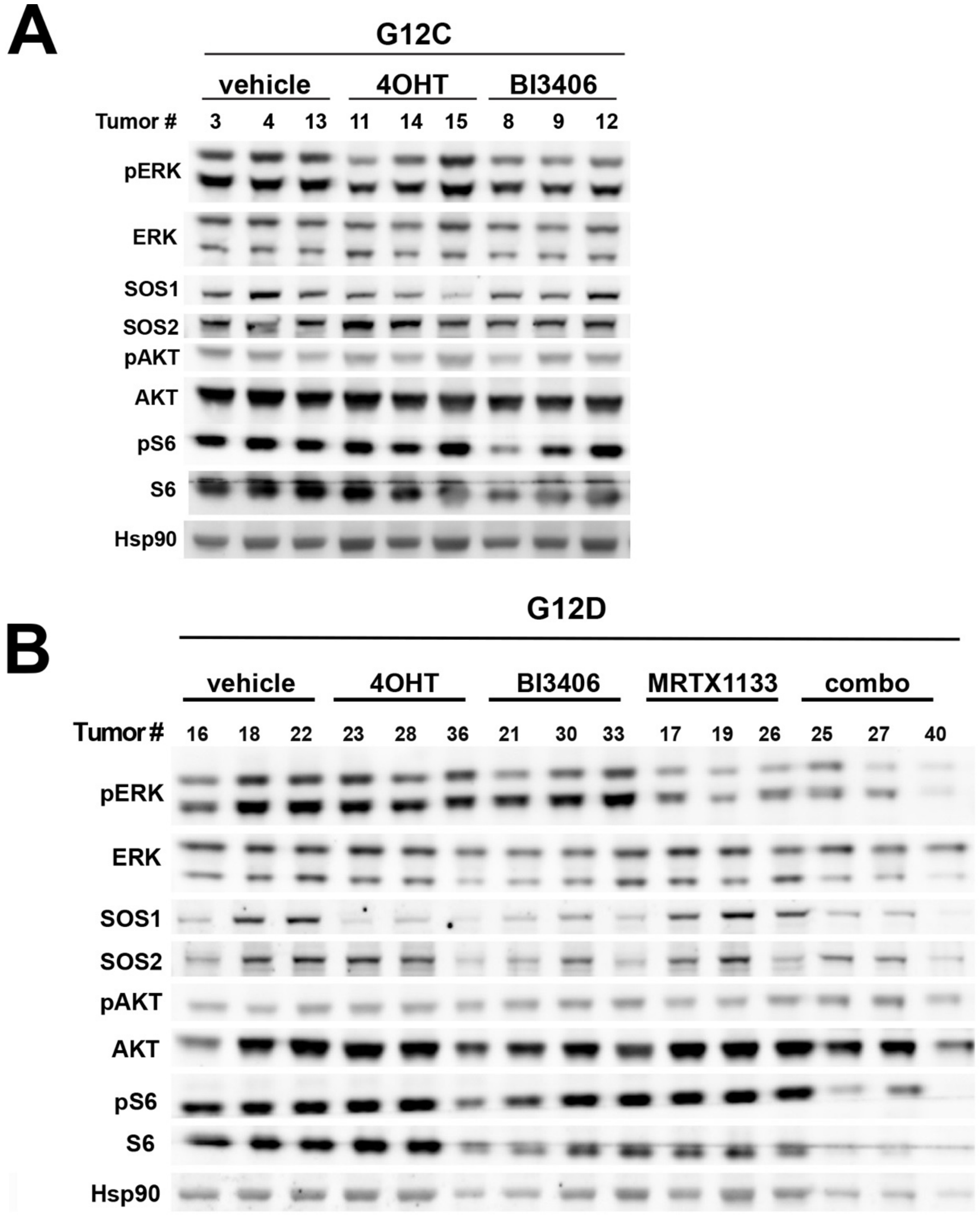


**Supplementary Figure S5. Influence of SOS1 ablation or inhibition in RAS/MAPK signal transmission in oncogenic KRAS tumors. (A-B)** Representative Western-blot images from KRAS$^{G12C}$ (A) and KRAS$^{G12D}$-mutated (B) tumor explants isolated from animals treated as indicated that were analyzed with the indicated antibodies.

### Supplementary Figure 6

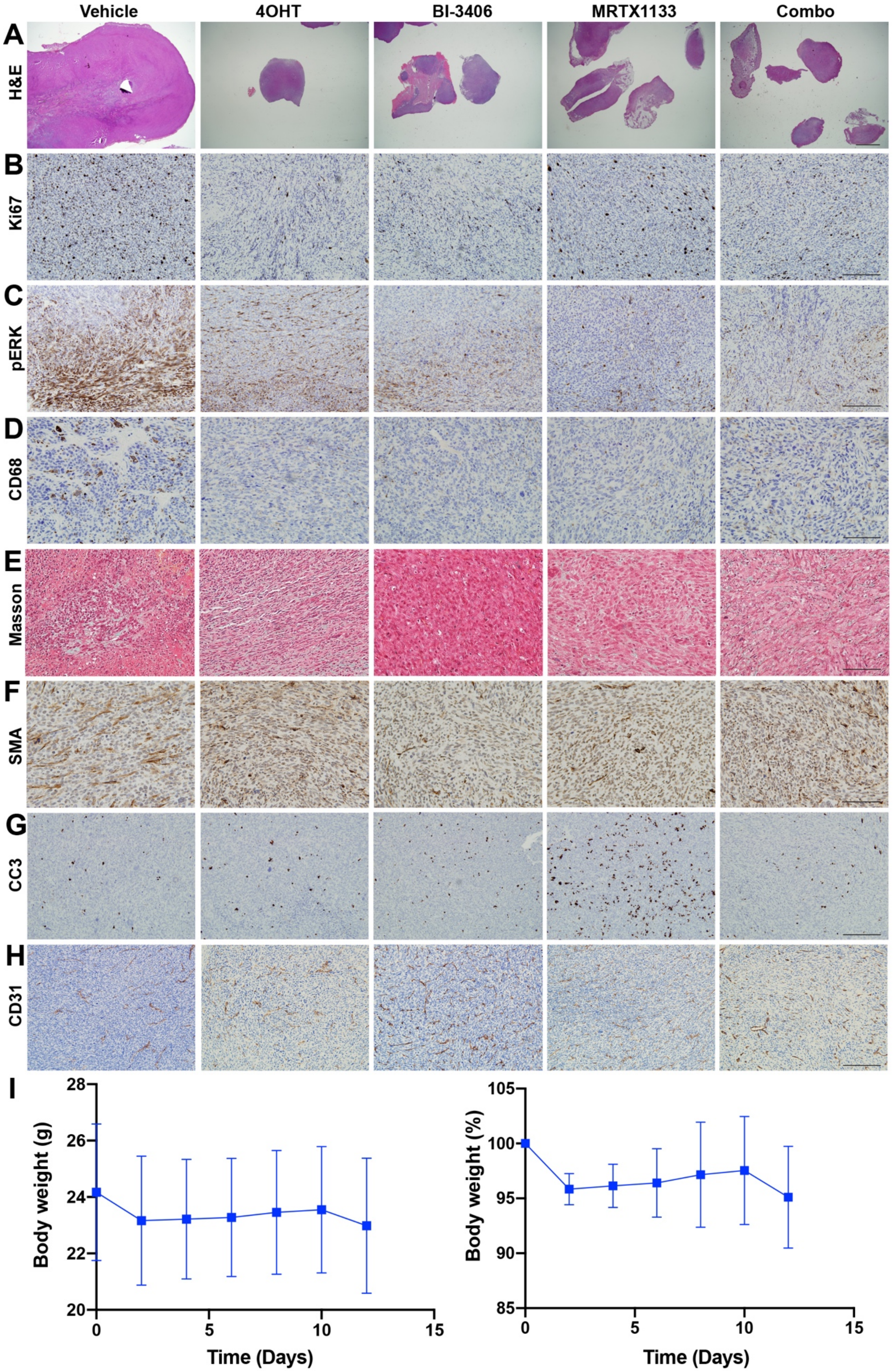

**Supplementary Fig. S6. Impact of concomitant targeting of SOS1 and KRAS$^{G12D}$ mutation on cell proliferation, cell death and tumor microenvironment in KRAS$^{G12D}$-mutated allografts.** Representative images of paraffin-embedded sections of isolated KRAS$^{G12D}$ mutated allografts from nude mice treated with vehicle, 4OHT (80 mg/kg), BI-3406 (100 mg/kg, once daily), MRTX1133 (30 mg/kg) or combo (BI-3406 plus MRTX1133) stained for H&E **(A)** and Masson's trichrome **(E)** or immunostained for Ki67 **(B)**, pERK **(C)**, CD68 **(D)**, SMA **(F)**, CC3 **(G)**, CD31 **(H)**. Scale bars: (A): 1 mm; (B, C, G, H): 100 µm; (D-F): 50 µm. **(I)** *In vivo* assessment of the effect of single or combined treatment with BI-3406 and MRTX1133 in KRAS$^{G12D}$-mutated allografts. Effects of MRTX1133 daily intraperitoneal administration for 12 days (30mg/kg/day) on body weight, total (left graph) or percentage (right graph) of nude mice (6-weeks old). n=5. Data shown as mean ± SD. CC3: cleaved caspase 3; SMA: smooth muscle actin.

**Supplementary Figure 7**

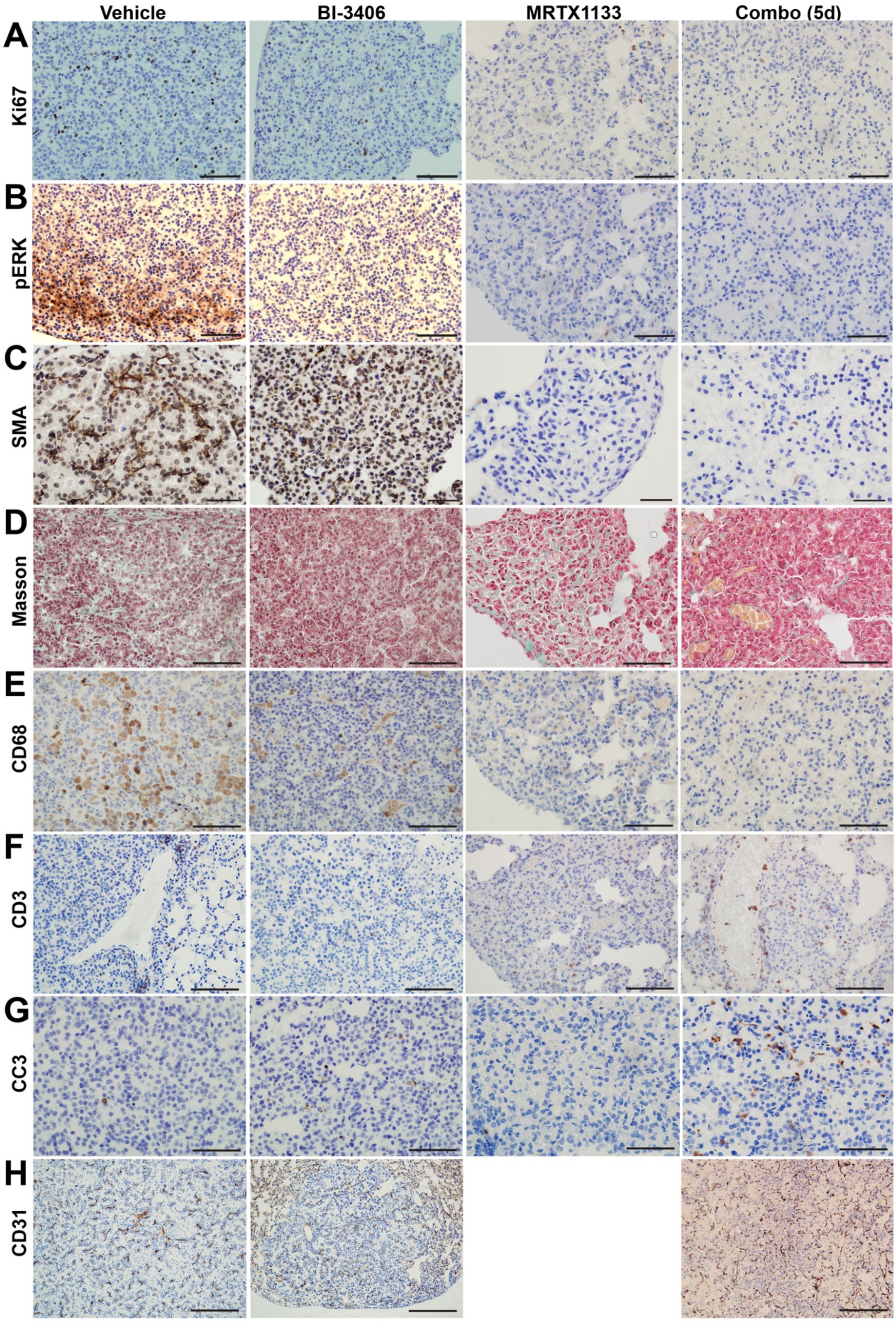


**Supplementary Figure S7**. **Effect of BI-3406 and MRTX1133 administration on the tumor microenvironment in KRAS$^{G12D}$-driven LUAD.** Representative images of paraffin-embedded sections from lungs of SOS1/2$^{WT}$/KRAS$^{G12D}$-mutated mice treated with the vehicle, BI-3406 (50 mg/kg, bid, for 26 days; starting when they were 3 months old), MRTX1133 (30 mg/kg for 17 days) or combo (BI-3406 50 mg/kg bid plus MRTX1133 30 mg/kg for 5 days) after immunostaining for Ki67 (**A**), pERK (**B**), SMA (**C**), CD68 (**E**), CD3 **(F)**, CC3 (G) and CD31 (H) or staining with Masson trichrome (**D**) as indicated. Please note that there is no panel showing CD31 immunostaining in the MRTX1133-treated group since the analyzed section did not include tumor cells and no other section was available. Scale bars: 50 μm. CC3: cleaved caspase 3; SMA: smooth muscle actin.

#### Supplementary Figure 8

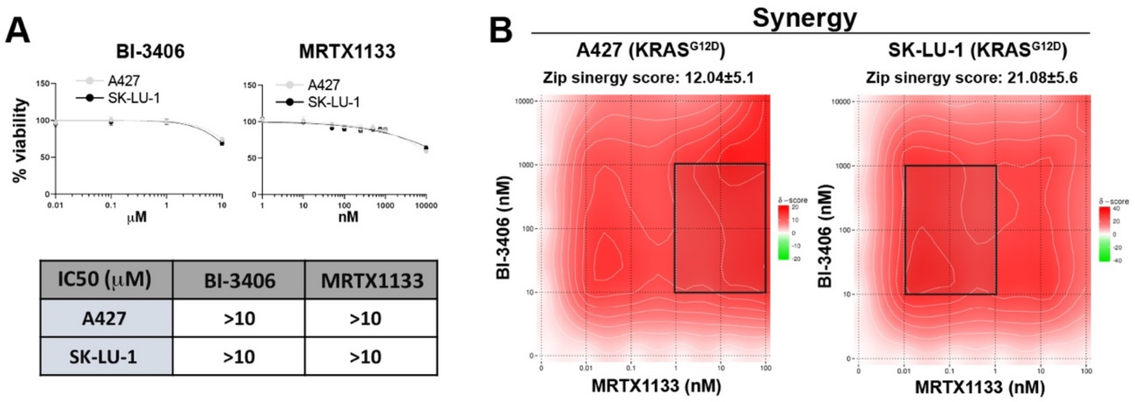


| IC50 (μM) | BI-3406 | MRTX1133 |
|---|---|---|
| A427 | >10 | >10 |
| SK-LU-1 | >10 | >10 |

**Supplementary Figure S8**. **Determination of sensitivity to BI-3406 and MRTX1133 in KRAS$^{G12D}$ mutant A247 and SK-LU-1 human LUAD cells. (A)** Curves of dose-response to BI-3406 and MRTX1133 in A427 or SK-LU-1 human LUAD KRAS$^{G12D}$ cell lines at the indicated concentrations. n=3 independent samples per experimental group. The table indicates that both BI-3406 and MRTX1133 show an IC50 higher than 10 μM when singly administered. **(B)** 2D synergy map representing the synergistic effects of BI-3406 and MRTX1133 combination at the indicated concentrations. Synergies were calculated based on the Zero Interaction Potency (ZIP) model. A Delta score higher than ten indicates that both drugs are likely synergistic. n=4 independent samples for A427 and n=5 independent samples for SK-LU-1 cells.

#### Supplementary Table 1

| | $SOS2^{KO}$ (Vehicle) | $SOS2^{KO}$ (BI-3406) |
|---|---|---|
| **Glc (mg/dl)** | 162±11 | 156±32 |
| **T-Chol (mg/dl)** | 138±14 | 134±23 |
| **BUN (mg/dl)** | 26±3.5 | 23.3±6 |
| **T-Bil** | 0.5±0.07 | 0.7±0.3* |
| **GOT (IU/liter)** | 66±28 | 103±37* |
| **GPT (IU/liter)** | 24±11 | 49±21* |
| **ALP (IU/liter)** | 292±34 | 310±71 |
| **CPK (IU/liter)** | 401±127 | 793±436 |
| **GGT (IU/liter)** | 10 | 10.7±4.1 |
| **T-Prot (g/dl)** | 6.44±0.5 | 6.15±0.6 |
| **Alb (g/dl)** | 3±0.15 | 2.8±0.23 |
| **Ca (mg/dl)** | 9.63±0.64 | 10.4±0.75 |
| **TG (mg/dl)** | 165±36 | 121±26** |
| **UA (mg/dl)** | 1.9±0.2 | 2.1±0.6 |
| **LDH (IU/liter)** | 738±363 | 1470±973 |

**Supplementary Table 1.** Blood serum parameters were assessed at day 26 after vehicle or BI-3406 oral administration (50 mg/kg, bid) in $SOS2^{KO}$ mice. Vehicle (n=7); BI-3406 (n=30). Data expressed as mean ± SD. * $p< 0.05$, ** $p< 0.01$ vs vehicle-treated $SOS2^{KO}$. Unpaired t-test was performed. **Alb**: Albumin; **ALP**: Alkaline Phosphatase; **BUN**: Blood urea nitrogen; **Ca**: Calcium; **CPK**: Creatine Phosphokinase; **GGT**: gamma-glutamyl transferase; **Glc**: Glucose; **GOT**: glutamic oxaloacetic transaminase; **GPT**: Glutamate-Pyruvate Transaminase; **LDH**: lactate dehydrogenase; **T-Bil**: Total bilirubin; **T-Chol**: Total Cholesterol; **TG**: Triglyceride; **T-Prot**: Total Protein; **UA**: **Uric acid**.

#### Supplementary Table 2

| Group | Organ | Feature | Severity | Observations |
|---|---|---|---|---|
| SOS1/2$^{WT}$ (TAM) | **Thymus** | Apoptosis/necrosis | 0 (8/8) | N/A |
| | **Spleen** | Apoptosis/necrosis | 0 (8/8) | N/A |
| | | Neutrophils influx | 0 (8/8) | |
| | | Thrombosis | 0 (8/8) | |
| | | Fibrosis | 0 (8/8)<br>1 (1/8) | |
| | **Liver** | Apoptosis/necrosis | 0 (8/8) | N/A |
| | | Inflammation | 0 (8/8) | |
| | | Fibrosis | 0 (8/8) | |
| | | Steatosis | 0 (8/8) | |
| | **Kidney** | Apoptosis/necrosis | 0 (8/8) | N/A |
| | | Inflammation | 0 (8/8) | |
| | | Tubular ectasia | 0 (8/8) | |
| | **Heart** | Apoptosis/necrosis | 0 (8/8) | N/A |
| | | Inflammation | 0 (8/8) | |
| | | Fibrosis | 0 (8/8) | |
| | **Lung** | Bronchitis | 0 (8/8) | N/A |
| | | Edema | 0 (8/8) | |
| | | Epithelial thickening | 0 (8/8) | |
| | | Apoptosis/necrosis | 0 (8/8) | |
| | | Metaplasia | 0 (8/8) | |
| | | Fibrosis | 0 (7/8)<br>1 (1/8) | |
| | | Neutrophilic inflammation | 0 (8/8) | |
| SOS1/2$^{DKO}$ (TAM) | **Thymus** | Apoptosis/necrosis | 1 (1/8)<br>2 (5/8)<br>3 (2/8) | Severe alteration of the histological structure |
| | **Spleen** | Apoptosis/necrosis | 1 (6/8)<br>2 (2/8) | Overall reduction of size without affectation in the histological architecture |
| | | Neutrophils influx | 0 (8/8) | |
| | | Thrombosis | 0 (8/8) | |
| | | Fibrosis | 0 (2/8)<br>1 (6/8) | |
| | **Liver** | Apoptosis/necrosis | 2 (7/8)<br>3 (1/8) | This organ shows an important rate of hepatocyte apoptosis |
| | | Inflammation | 0 (8/8) | |
| | | Fibrosis | 0 (8/8) | |
| | | Steatosis | 0 (8/8) | |
| | **Kidney** | Apoptosis/necrosis | 0 (8/8) | N/A |
| | | Inflammation | 0 (8/8) | |
| | | Tubular ectasia | 0 (8/8) | |
| | **Heart** | Apoptosis/necrosis | 0 (8/8) | N/A |
| | | Inflammation | 0 (8/8) | |
| | | Fibrosis | 0 (8/8) | |
| | **Lung** | Bronchitis | 0 (8/8) | N/A |
| | | Edema | 0 (8/8) | |
| | | Epithelial thickening | 0 (7/8)<br>1 (1/8) | |
| | | Apoptosis/necrosis | 0 (8/8) | |
| | | Metaplasia | 0 (8/8) | |
| | | Fibrosis | 0 (6/8)<br>1 (2/8) | |
| | | Neutrophilic inflammation | 0 (8/8) | |
| SOS1/2$^{WT}$ (BI-3406) | **Thymus** | Apoptosis/necrosis | 0 (10/12)<br>1 (2/12) | N/A |
| | **Spleen** | Apoptosis/necrosis | 0 (11/12)<br>1 (1/12) | N/A |
| | | Neutrophils influx | 0 (12/12) | |

| | | | | |
|---|---|---|---|---|
| | | Thrombosis | 0 (12/12) | |
| | | Fibrosis | 0 (11/12)<br>1 (1/12) | |
| | **Liver** | Apoptosis/necrosis | 0 (11/12)<br>1 (1/12) | N/A |
| | | Inflammation | 0 (12/12) | |
| | | Fibrosis | 0 (12/12) | |
| | | Steatosis | 0 (12/12) | |
| | **Kidney** | Apoptosis/necrosis | 0 (12/12) | N/A |
| | | Inflammation | 0 (12/12) | |
| | | Tubular ectasia | 0 (12/12) | |
| | **Heart** | Apoptosis/necrosis | 0 (12/12) | N/A |
| | | Inflammation | 0 (12/12) | |
| | | Fibrosis | 0 (12/12) | |
| | **Lung** | Bronchitis | 0 (12/12) | N/A |
| | | Edema | 0 (12/12) | |
| | | Epithelial thickening | 0 (12/12) | |
| | | Apoptosis/necrosis | 0 (12/12) | |
| | | Metaplasia | 0 (12/12) | |
| | | Fibrosis | 0 (12/12) | |
| | | Neutrophilic inflammation | 0 (12/12) | |
| **SOS2$^{KO}$ (BI-3406)** | **Thymus** | Apoptosis/necrosis | 0 (9/12)<br>1 (3/12) | N/A |
| | **Spleen** | Apoptosis/necrosis | 0 (11/12)<br>1 (1/12) | N/A |
| | | Neutrophils influx | 0 (12/12) | |
| | | Thrombosis | 0 (12/12) | |
| | | Fibrosis | 0 (9/12)<br>1 (3/12) | |
| | **Liver** | Apoptosis/necrosis | 0 (8/12)<br>1 (4/12) | Clusters of hepatocytes with smaller and darker nuclei and retracted cytoplasm (apoptotic features) were intermittently observed |
| | | Inflammation | 0 (12/12) | |
| | | Fibrosis | 0 (12/12) | |
| | | Steatosis | 0 (12/12) | |
| | **Kidney** | Apoptosis/necrosis | 0 (12/12) | Some epithelial cells covering the surface of the proximal and distal tubules showed cytoplasmic vacuoles |
| | | Inflammation | 0 (12/12) | |
| | | Tubular ectasia | 0 (12/12) | |
| | **Heart** | Apoptosis/necrosis | 0 (12/12) | N/A |
| | | Inflammation | 0 (12/12) | |
| | | Fibrosis | 0 (12/12) | |
| | **Lung** | Bronchitis | 0 (12/12) | N/A |
| | | Edema | 0 (12/12) | |
| | | Epithelial thickening | 0 (11/12)<br>1 (1/12) | |
| | | Apoptosis/necrosis | 0 (12/12) | |
| | | Metaplasia | 0 (12/12) | |
| | | Fibrosis | 0 (9/12)<br>1 (3/12) | |
| | | Neutrophilic inflammation | 0 (12/12) | |
| **SOS1/2$^{WT}$ (vehicle)** | **Thymus** | Apoptosis/necrosis | 0 (7/7) | N/A |
| | **Spleen** | Apoptosis/necrosis | 0 (7/7) | N/A |
| | | Neutrophils influx | 0 (7/7) | |
| | | Thrombosis | 0 (7/7) | |
| | | Fibrosis | 0 (7/7) | |
| | **Liver** | Apoptosis/necrosis | 0 (7/7) | N/A |
| | | Inflammation | 0 (7/7) | |
| | | Fibrosis | 0 (7/7) | |
| | | Steatosis | 0 (7/7) | |
| | **Kidney** | Apoptosis/necrosis | 0 (7/7) | N/A |
| | | Inflammation | 0 (7/7) | |
| | | Tubular ectasia | 0 (7/7) | |

| | | | | |
|---|---|---|---|---|
| | **Heart** | Apoptosis/necrosis | 0 (7/7) | N/A |
| | | Inflammation | 0 (7/7) | |
| | | Fibrosis | 0 (7/7) | |
| | **Lung** | Bronchitis | 0 (7/7) | N/A |
| | | Edema | 0 (7/7) | |
| | | Epithelial thickening | 0 (7/7) | |
| | | Apoptosis/necrosis | 0 (7/7) | |
| | | Metaplasia | 0 (7/7) | |
| | | Fibrosis | 0 (6/7)<br>1 (1/7) | |
| | | Neutrophilic inflammation | 0 (7/7) | |
| **SOS2$^{KO}$ (vehicle)** | **Thymus** | Apoptosis/necrosis | 0 (7/7) | N/A |
| | **Spleen** | Apoptosis/necrosis | 0 (7/7) | N/A |
| | | Neutrophils influx | 0 (7/7) | |
| | | Thrombosis | 0 (7/7) | |
| | | Fibrosis | 0 (5/7)<br>1 (2/7) | |
| | **Liver** | Apoptosis/necrosis | 0 (7/7) | N/A |
| | | Inflammation | 0 (7/7) | |
| | | Fibrosis | 0 (7/7) | |
| | | Steatosis | 0 (7/7) | |
| | **Kidney** | Apoptosis/necrosis | 0 (7/7) | N/A |
| | | Inflammation | 0 (7/7) | |
| | | Tubular ectasia | 0 (7/7) | |
| | **Heart** | Apoptosis/necrosis | 0 (7/7) | N/A |
| | | Inflammation | 0 (7/7) | |
| | | Fibrosis | 0 (7/7) | |
| | **Lung** | Bronchitis | 0 (7/7) | N/A |
| | | Edema | 0 (7/7) | |
| | | Epithelial thickening | 0 (7/7) | |
| | | Apoptosis/necrosis | 0 (7/7) | |
| | | Metaplasia | 0 (7/7) | |
| | | Fibrosis | 0 (6/7)<br>1 (1/7) | |
| | | Neutrophilic inflammation | 0 (7/7) | |

**Supplementary Table 2.** Histological scoring. TAM-treated SOS1/2$^{WT}$ (n=8); TAM-treated SOS1/2$^{DKO}$ (n=8); BI-3406-treated SOS1/2$^{WT}$ (n=12); BI-3406-treated SOS2$^{KO}$ (n=12); vehicle-treated SOS1/2$^{WT}$ (n=7); vehicle-treated SOS2$^{KO}$ (n=7).
